# Fast tracking native mass spectrometry: Skipping over buffer exchange

**DOI:** 10.1101/2025.02.22.639503

**Authors:** Alice Frederike Rosa Grün, Fatema-Aqila Said, Kira Schamoni-Kast, Tomislav Damjanović, Assel Berikkara, Jonas Schröder, Fenja Kleine Brockmann, Tobias Lichtenberg, Jens Bosse, Charlotte Uetrecht

## Abstract

Obtaining sufficient amounts of pure protein for downstream applications such as native mass spectrometry (nMS) is often challenging, especially when expression yields are low or proteins are unstable. In these cases, the commonly required buffer-exchange step is a major bottleneck, as it often leads to substantial protein loss and compromises biophysical characterization.

These challenges are exacerbated in insect or eukaryotic expression systems, where protein yields are typically lower than in bacteria, making protein loss during purification particularly detrimental. Standard lysis and purification buffers contain non-volatile components such as Tris, phosphate, HEPES and sodium chloride, which form adducts during electrospray ionization (ESI) interfering with the signal and therefore must be removed prior to nMS. To address protein loss associated with this mandatory buffer-exchange, we evaluated an affinity-purification workflow, in which non-volatile salts are excluded throughout purification and proteins are directly eluted into nMS-compatible ammonium acetate– based buffers. This approach eliminates the need for a separate buffer exchange step and enables rapid nMS analysis immediately after affinity purification. We show that common eluents used in His- and Strep-based affinity purification, such as imidazole, biotin, and desthiobiotin, are well tolerated at relevant concentrations, allowing acquisition of high-quality spectra suitable for determining protein stoichiometry and for monitoring enzymatic or assembly processes.

Together, this fast-track affinity workflow increases protein recovery, shortens sample preparation and complements online exchange protocols, which are less suited for monitoring processes. It hence expands the applicability of nMS to proteins and protein complexes that are difficult to obtain in sufficient quantity using conventional purification and buffer exchange strategies.

**TOC Graphic:** 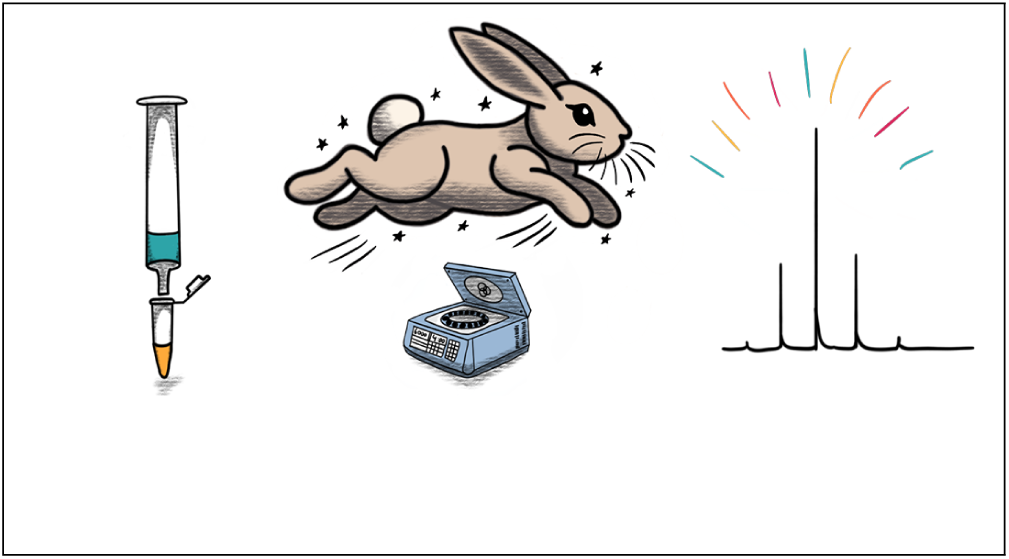

## Introduction

Protein expression in insect or mammalian cells generally results in lower yields compared to bacterial expression. Moreover, eukaryotic expression is typically chosen when proteins show impaired folding upon bacterial expression or when post-translational modifications (PTMs) are relevant to function. In turn, proteins that do not fold independently often have reduced stability limiting their shelf-life. Additional purification steps can hence complicate their characterization by native mass spectrometry (nMS).^1^ These challenges make sample handling a critical factor for maintaining protein integrity and obtaining high-quality spectra. To avoid unnecessary protein loss during purification, the “direct MS” approach was developed. Well-expressed proteins are directly sprayed into the mass spectrometer after lysing cells (intracellular proteins) or diluting the cell culture supernatant (secreted proteins) in high concentrations of ammonium acetate (AmAc).^2,3^This “direct MS” approach minimizes sample handling and has proven highly effective for abundant proteins, but has limited applicability for proteins expressed at low levels requiring enrichment or purification prior to analysis. A recent approach, termed SNAP-MS, addresses this issue by using an elution-free affinity purification protocol that employs dissolvable hydrogel microbeads.^4^ The protocol has been employed to purify and characterize low-abundance intact proteins, including those extracted from blood samples. While SNAP-MS provides an important advance for low-input native MS workflows, it relies on specialized bead materials and a dedicated bead-dissolution strategy, with a reported sample-to-data time of approximately 2 h.^4^ Thus, for unstable or time-sensitive recombinant proteins, even faster workflows with minimal post-purification handling remain desirable. Several online chromatographic approaches have also been developed to streamline nMS workflows, including online affinity-LC-native MS for antibodies and other targets, online size exclusion chromatography (SEC)-native MS for soluble and membrane proteins, and related integrated platforms.^5–7^ These methods provide excellent separation and compatibility with nMS, but they often dilute the sample, and offer limited flexibility for downstream biochemical and biophysical characterization prior to nMS. Therefore, workflows that minimize sample handling while maintaining nMS compatibility, avoiding sample dilution, and preserving peak resolution remain highly desirable. Standard workflows for purification of proteins from expression systems often utilize affinity chromatography. Two commonly used affinity-tags are His-tags and StrepII-tags.^8^ His-tags bind to Ni(II)^9^ and are eluted by imidazole in concentrations ranging from 100 to 500 mM.^10^ This system is cost-efficient but often requires an additional SEC due to low specificity. Ni(II) is complexed by histidine stretches, which can occur naturally in proteins, and by metal-binding proteins from cell culture media, leading to unspecific binding.^9,11,12^ The StrepII-tag (as well as related tags) mimics the interaction between streptavidin and biotin, which has one of the highest binding affinities found in nature.^13^ As biotinylation is a naturally uncommon protein modification, unspecific binding is significantly reduced,^14^ resulting in very pure protein samples even in eukaryotic expression.^15^ Here, elution buffers usually contain 50 mM biotin or 2.5 mM desthiobiotin, with the latter having slightly lower affinity, allowing for easier column recovery.^16^ In both cases, protein purification is compatible with most standard buffers (e.g. Tris-, HEPES-, or phosphate) supplemented with sodium chloride, which stabilizes the proteins, but complicates nMS. During protein ionization via nano-electrospray ionization (nESI), non-volatile salts like sodium chloride and the buffer molecules form adducts on the species of interest and salt clusters. These adducts spread out mass signals and decrease signal intensity up to even completely masking protein peaks in the spectra.^17–19^ Although it is possible to reduce salt adducts using e.g. submicron emitters enabling nMS from regular buffer solutions, such studies have been mostly employed on smaller standard proteins.^18,20,21^ This revealed however that buffer composition is relevant for observed conformation.^21,22^ Nevertheless, buffer exchange is still an integral part of sample preparation for nMS-based structural proteomics in many cases. Buffer exchange is often performed via spin filters, spin columns, SEC, or dialysis cartridges. ^23–28^ Unfortunately, proteins often bind irreversibly to the different matrices or even precipitate, resulting in significant protein loss. Moreover, buffer exchange increases handling time between cell lysis and nMS^23^ which can be detrimental for structurally labile proteins or protein complexes. As described above, previous efforts to reduce salt adducts highlight the clear advantage of avoiding buffer exchange altogether.^6,23,29^

Here, we systematically evaluated affinity-purification workflows for His- and StrepII-tags, which exclude non-volatile salts during lysis, purification and, importantly, during elution. Instead, we used volatile AmAc-based buffer surrogates that are compatible with nMS at appropriate concentrations and pH levels to avoid subsequent buffer exchange. To evaluate if commonly used eluent concentrations of imidazole, biotin, and desthiobiotin were compatible with obtaining charge state resolved spectra, we employed alcohol dehydrogenase (ADH) as standard protein showing that imidazole and desthiobiotin are suitable with the latter performing best. We then tested the protocol with a variety of viral proteins: *E. coli* expressed His- and Strep-tagged SARS-CoV-2 non-structural protein 7-11 (nsp7-11) polyprotein, HEK293T cell expressed SARS-CoV-2 nucleocapsid (N) protein, His- and Strep-tagged SARS-CoV-2 nsp12 expressed in both *E. coli* and HEK293T cells, and different herpesvirus proteins expressed in mammalian (KSHV pORF68) and insect cells (HSV-1 pUL32 and the terminase complex out of pUL15, pUL28, pUL33). Importantly, the eluents were also compatible with the processing of nsp7-11 by its cognate main protease M^pro^, indicating that the workflow is indeed compatible with subsequent biophysical characterization including enzymatic processing and subsequent complex formation. Since time from cell lysis over purification to nMS is below 2 h, we coined this approach “fast-track sample preparation protocol”.

## Materials and Methods

### ADH sample preparation

6 mg of ADH (37 kDa, Alcohol Dehydrogenase from *Saccharomyces cerevisiae*, A7011-75KU, Sigma Aldrich) was dissolved in 100 µL MilliQ water, and 50 µL was loaded onto a Biospin column (Biospin mini columns, 6000 MWCO, Bio-Rad) that was equilibrated with 150 mM ammonium acetate (AmAc) (99.99 % purity, 431311, Sigma Aldrich) at pH 8. The spin column was centrifuged at 1000 *x g* at 4 *^◦^*C. ADH in AmAc was eluted and diluted to a concentration of 7.5 µM and 2.5 µM. From this sample, aliquots were taken and spiked with different concentrations of imidazole (A10221.22, Thermo Fisher Scientific), biotin (2-1016- 002, IBA) and desthiobiotin (2-1000-002, IBA).

### Expression, purification and processing of nsp7-11 in *E. coli*

The SARS-CoV-2 (original Wuhan-Hu-1 variant) polyprotein nsp7-11-His_6_ (NCBI LOCUS NC_045512) was expressed as previously described.^30^ Briefly, the polyprotein was expressed in BL21 (DE3) cells overnight at 37 *^◦^*C. Cells were spun down by centrifugation and lysed in lysis buffer (300 mM AmAc, 1 mM dithiothreitol (DTT) (D0632, Sigma Aldrich) and 0.2 mg mL*^−^*^1^ lysozyme (L6876, Sigma Aldrich) at pH 8) via sonication. Cell debris was then removed by centrifugation, afterward, affinity purification of the polyprotein was performed on PureCube 100 Ni-INDIGO Agarose resin (75103, Cube Biotech). The resin was washed with 10 bed volumes (BV) of the wash buffer (300 mM AmAc, 1 mM DTT at pH 8) and elution was done with 0.5 BV elution buffer (300 mM AmAc, 1 mM DTT, 300 mM imidazole at pH 8). The expression and lysis of Strep -nsp7-11 was performed identical to nsp7-11-His_6_. Affinity purification utilized streptavidin resins (Strep-Tactin^®^ Sepharose^®^ column, 1 mL, 2-1202-001, IBA). The elution buffer contained 2.5 mM desthiobiotin instead of imidazole. To probe protease cleavage, a sample containing 15 µM polyprotein and 3 µM M^pro31^ was incubated at 4 *^◦^*C overnight and then directly sprayed into the MS.

### Expression and purification of nsp12 in mammalian cells and in *E. coli*

SARS-CoV-2 nsp12 with a Twin-Strep-II tag (141378, Addgene) was expressed in HEK293 human embryonic kidney cells. HEK293 cells were cultured in DMEM supplemented with 10% fetal bovine serum (FBS) and 1% penicillin/streptomycin. Transfection was performed using the PEI STAR transfection reagent (7854, Bio-Techne GmbH) with a plasmid encoding nsp12-StrepII. Transfected cells were selected using 2.5 μg/mL puromycin and subsequently subcultured in 250 mL Freestyle medium (12338018, Thermo Fisher). Cells were harvested at a density of 3–5 × 10⁶ cells/mL and stored at −70 °C until further use.

For affinity purification, cells were lysed in 500 mM AmAc, pH 8.0, supplemented with 1 mM tris(2-carboxyethyl)phosphine (TCEP) (T2556, Thermo Fischer). Washing was performed in the same buffer system. The lysate was applied to Strep-Tactin resin with a bed volume of 250 μL (2-1202-001, IBA). Recombinant nsp12 was eluted with wash buffer supplemented with 2.5 mM desthiobiotin (2-1000-005, IBA).

SARS-CoV-2 nsp12 carrying an N-terminal Strep-II tag was also expressed in *E. coli* BL21(DE3) cells as described for nsp7-11. The same fast-track protocol was used as for nsp12 expressed in mammalian cell line. Lysozyme was added to the lysis buffer (500 mM AmAc, 2 mM DTT (D0632, Sigma Aldrich), pH 8) at a final concentration of 0.2 mg/mL (L6876, Sigma-Aldrich). Affinity purification was performed using Strep-Tactin resin with a bed volume of 250 μL (2-1202-001, IBA). The elution of recombinant protein was performed in wash buffer with 2.5 mM desthiobiotin (2-1000-005, IBA).

### Expression and purification of the SARS-CoV2 N-protein in mammalian cells

For the standard purification protocol, stable suspension-adapted HEK293T cells expressing the SARS-CoV-2 Nucleocapsid (N) protein with a C-terminal Twin-Strep-tag (gift from Nevan Krogan; Addgene plasmid #141391)^32^ were harvested and resuspended in two pellet volumes of lysis buffer (150mM NaCl, 50 mM Tris HCL ph8, 1% P-nonidet p-40, 0.5% sodium desoxycholate, 0.5 mM EDTA, 0.1% SDS) supplemented with protease inhibitors (4693132001, Roche). Cells were disrupted by sonication and lysate was cleared by centrifugation (20,000 × g). The supernatant was loaded onto a Strep-Tactin® gravity flow column (IBA Lifesciences) equilibrated with 5 BV of Buffer W. The column was washed with 10 BV of Buffer W containing 10 mM ATP and 20 mM MgCl₂, followed by a second wash step with 10 BV of standard Buffer W. The target protein was eluted in six 0.5 mL fractions using Buffer E (IBA Lifesciences).

For the fast-track approach, the same cells were harvested and resuspended in two pellet volumes of 500 mM AmAc (pH 8.0) supplemented with protease inhibitors. Cells were disrupted by sonication. The lysate was cleared by centrifugation (20,000 × g) and sterile-filtered. The supernatant was loaded onto a Strep-Tactin® gravity flow column (IBA Lifesciences) equilibrated with 5 BV of 500 mM AmAc. To remove impurities, the column was washed with 10 BV of 500 mM AmAc containing 10 mM ATP and 20 mM MgCl₂, followed by a second wash step with 10 BV of 500 mM AmAc alone. The target protein was eluted in six 0.5 mL fractions using 500 mM AmAc supplemented with 2.5 mM desthiobiotin.

### Expression and purification of the terminase complex in insect cells

The terminase subunits (pUL15, pUL28 and pUL33) of herpes simplex virus 1 (HSV-1) were expressed in 2x25 mL of ExpiSf9™ (A35243, Thermo Fisher) suspension culture by infecting the cells with two baculoviruses. These were generated from Bac-to-Bac protocol from a pFastBac1 His-UL15 plasmid, and a pFastBacDual plasmid containing UL28 and UL33. They were harvested 3 dpi by centrifugation (5 min, 4 *^◦^*C, 300 x*g*) and the pellet washed in PBS once and then stored at *-*70 *^◦^*C. The pellet was then thawed at 4 *^◦^*C in 10 mL lysis buffer (1 M AmAc, 1 mM DTT, and EDTA-free protease inhibitors (4693132001, Roche), pH 7.5). The lysate was then sonicated and applied to a gravity column containing PureCube 100 Ni-INDIGO Agarose resin. The beads were washed with washing buffer A (300 mM AmAc, 1 mM DTT, pH 7.5) for 20 BV. Afterward, the column was washed with washing buffer B (300 mM AmAc, 1 mM DTT, 35 mM imidazole, pH 7.5) until the concentration of the solution reached zero mg/mL. The terminase complex was then eluted in steps of 300 µL elution buffer (300 mM AmAc, 1 mM DTT, 350 mM imidazole, pH 7.5) for 5 BV. For the terminase, triplicate measurements of the same purified sample were obtained.

### Expression and purification of pUL32 in insect cells

The HSV-1 terminase-associated protein pUL32 was expressed in ExpiSf9™ cells. A 100 mL cell suspension was infected with a baculovirus containing the UL32 gene at an MOI of 3. The baculovirus was generated via Bac-to-Bac protocol from a pFastBac1 StrepII-UL32 plasmid. The cells were harvested at 3 dpi by centrifugation (5 min, 4 *^◦^*C, 300 x*g*). The pellet was then washed once in PBS and stored at *-*70 *^◦^*C for a few days. The pellet was thawed in three times the pellet volume of ice-cold lysis buffer (1 M AmAc, 1 mM DTT, 0.1 % CHAPS (1479.1, Carl Roth) and EDTA-free protease inhibitors, pH 8). The cells were lysed via sonication on ice. The lysate was added to a Strep-Tactin Sepharose column. The column was then washed with 5 BV of washing buffer (400 mM AmAc, 1 mM DTT, and 0.1 % CHAPS, pH 8). Six times 0.5 BV of elution buffer (washing buffer with 2.5 mM desthiobiotin) were used to displace the proteins. The samples were then mixed with a 2 µM of a purified terminase sample (1:1) before nMS measurement. For pUL32 triplicate measurements of the same purified sample were performed.

### Expression and purification of pORF68 in mammalian cells

The expression of human herpesvirus 8 (KSHV HHV-8) pORF68 was performed in 100 mL FreeStyle™ 293-F-cells (R79007, Thermo Fisher). The cells were transfected with 10 µg of a mNeonGreen and 115 µg of an ORF68 containing plasmid (pcDNA4/TO-2xStrep-ORF68 was a gift from Britt Glaunsinger (Addgene plasmid #162625; http://n2t.net/addgene:162625; RRID: Addgene_162625). They were harvested by centrifugation after 3 dpi, and frozen at*−*70 *^◦^*C. Otherwise, the purification of pORF68 was identical to that of pUL32, but no terminase was added prior measurement. For pORF68 only one measurement was obtained.

### Native mass spectrometry

In order to transfer the sample into the mass spectrometer, nESI capillaries were pulled on a micropipette puller (P-1000, Sutter Instruments). A two-step method to heat up a squared-box filament (2.5 mm x 2.5 mm) was used for pulling the borosilicate capillaries (1.2 mm outer and 0.68 mm inner diameter, Kwik-Fil 1B120F-4, World Precision Instruments). The glass capillaries were then gold-coated in a sputter coater (CCU-010, Safematic, 5.0 *×* 10*^−^*^2^ mbar, 30.0 mA, 120 s, three runs to vacuum limit 3.0 *×* 10*^−^*^2^ mbar argon). Three technical replicates were obtained per sample unless indicated otherwise.

Measurements were performed on two systems. A Q-TOF 2 (Waters, Manchester, UK) modified for high masses (MS Vision, Almere, the Netherlands)^33^ and mass calibrated with CsI (25 mg mL*^−^*^1^); and on a QExactive UHMR Orbitrap (Thermo Fisher) with a resolution of 6250 at 400 *m*/*z* throughout. Positive ion mode was used on both systems, sample specific instrument settings are provided in table 1 (Q-TOF 2) and table 2 (QE UHMR).

**Table 1:** Q-TOF2 settings.

| Parameter | ADH | nsp7-11 |
| --- | --- | --- |
| capillary voltage (eV) | 1.35 | 1.35 |
| cone voltage | 150 | 150 |
| source press. (mbar) | 10 | 10 |
| cone temp. (°C) | 80 | 80 |
| collisional gas | Argon | Argon |
| collisional volt. (CV) | 10-400 | 20-200 |
| collisional gas press.<br>(mbar) | 1.8x10<sup>-2</sup> | 1.8x10<sup>-2</sup> |
| acquisition window<br>(*m/z*) | 100-30000 | 100-20000 |

**Table 2:** QExactive UHMR orbitrap settings.

| Parameter | ADH | nsp7-11 | nsp12 | N-protein | terminase | pUL32 | pORF68 |
| --- | --- | --- | --- | --- | --- | --- | --- |
| capillary<br>voltage (eV) | 1.35 | 1.30 | 1.3 | 1.7 | 1.5 | 1.5 | 1.5 |
| capillary<br>temp. (°C) | 80 | 150 | 150 | 100 | 250 | 250 | 250 |
| Inj. Fl. RF<br>Ampl. | 300 | 300 | 700 | 700 | 700 | 700 | 700 |
| Bent. Fl. RF<br>Ampl. | 940 | 940 | 940 | 940 | 940 | 940 | 940 |
| Trans MP | 900 | 900 | 900 | 900 | 900 | 900 | 900 |
| HCD-cell RF<br>Ampl. | 900 | 900 | 900 | 900 | 900 | 900 | 900 |
| collisional<br>volt. (eV) | 0-300 | 0-200 | 0-50 | 55 | 0-200 | 0-200 | 0-200 |
| In-source<br>CID (eV) | 0 | 0 | 0-20 | 25 | 0 | 0 | 0 |
| Detector op-<br>tim. | Low *m/z* | Low *m/z* | Low *m/z* | High *m/z* | High *m/z* | Low *m/z* | High *m/z* |
| Trap. gas<br>pressure set-<br>ting | 6 | 6 | 5 | 5 | 5 | 7 | 5 |
| C-Trap<br>Ampl. | 2950 | 2750 | 2950 | 2950 | 2950 | 2950 | 2950 |

### Data analysis

Mass spectra obtained from the Q-TOF 2 were analyzed with MassLynx™ (ver. 4.1, Waters, Manchester, UK) and spectra obtained with the QExactive UHMR were analyzed with Xcalibur (ver. 4.2.47, Thermo Fisher). For well-resolved spectra, initial mass determination was performed using UniDec. For poorly resolved or imidazole-containing spectra, where automated peak picking and deconvolution were less reliable, Massign^34^ and a custom Python script were used for manual peak assignment, mass calculation, and full-width half-maximum (FWHM) determination. The script extracted defined m/z regions, identified charge-state peaks, and calculated masses from the corresponding m/z values and charge states. Mass and FWHM values from triplicate measurements were then averaged, and the standard error of the mean (SEM) was reported. The spectra were then exported into Adobe Illustrator (ver. 19.5.0.84, Adobe), for figure preparation. A high-quality spectrum is defined here as having charge-state resolved spectra, i.e. it exhibiting peak separation in which adjacent peaks are distinct down to at least half their maximum height.

## Results and discussion

### Influence of high eluent concentrations on spectral quality

To assess if protein complexes could be identified via nMS in the presence of high concentrations of imidazole, biotin or desthiobiotin, we first assessed their impact using commercially available, purified, tetrameric ADH. Data was obtained on two mass spectrometers, which are known to have substantially different desolvation/declustering capacity in the source region, with a Q-TOF 2 (Figure 1) representing “soft conditions” as also encountered on Synapt setups and with a QExactive UHMR (Figure 2) representing a “hot” source. Even under soft conditions on the Q-TOF2 (Figure 1), mass spectra with commonly used eluent concentrations such as 20 mM biotin, 2.5 mM desthiobiotin, or 300 mM imidazole still produce high-quality, charge-state resolved spectra of tetrameric ADH, whereby a high-quality spectrum is defined by essentially baseline separated charge states. Good spectra were obtained at medium collision voltages (CV) for all tested eluents, allowing for further increase of CV for collision-induced dissociation (CID) if needed. At low CV, spectra showed cluster formation for biotin and desthiobiotin, which were easily disrupted at increased CV (see Figure S1, S2 and S3). After further declustering of desthiobiotin and biotin, the resulting spectra looked similar to ADH without eluents, and charge state distributions were nearly identical. Solely for imidazole, peak broadening was observed compared to ADH spectra without eluent. Even though His-tag purification requires a large amount of imidazole, the CV needed to obtain a useful spectrum was comparable to the other two eluents. As opposed to biotin and desthiobiotin, imidazole is a charge-reducing agent, which leads to peaks at higher *m/z*.^35^ On the Q-TOF 2, higher amounts of eluent resulted in no longer resolvable peaks independent of the CV employed. The upper limit for biotin was 20 mM, which implies that at least a 2.5-fold dilution of the purified sample would be required after elution with the standard eluent concentration of 50 mM. This also applies to imidazole when concentrations higher than 300 mM are required. Strikingly, the upper concentration limit for desthiobiotin was not even reached at 25 mM, far beyond requirements for purification protocols that range from 2.5 mM^16,36^ to 5 mM^37^. Biotin and desthiobiotin also increased the number of detectable charge states compared to pure or imidazole-containing ADH solutions. This can be advantageous when deconvoluting the spectra, since more peaks increase the confidence of the calculated mass. Notably, the QExactive UHMR tolerated even higher amounts of eluent, and obtaining peak resolution was unproblematic at all tested eluent concentrations (see Figure 2). Again, higher than necessary concentrations could be used for desthiobiotin, but now mass spectra with 50 mM biotin or 500 mM imidazole were also resolved (Figure 2, S3).

**Figure 1:**
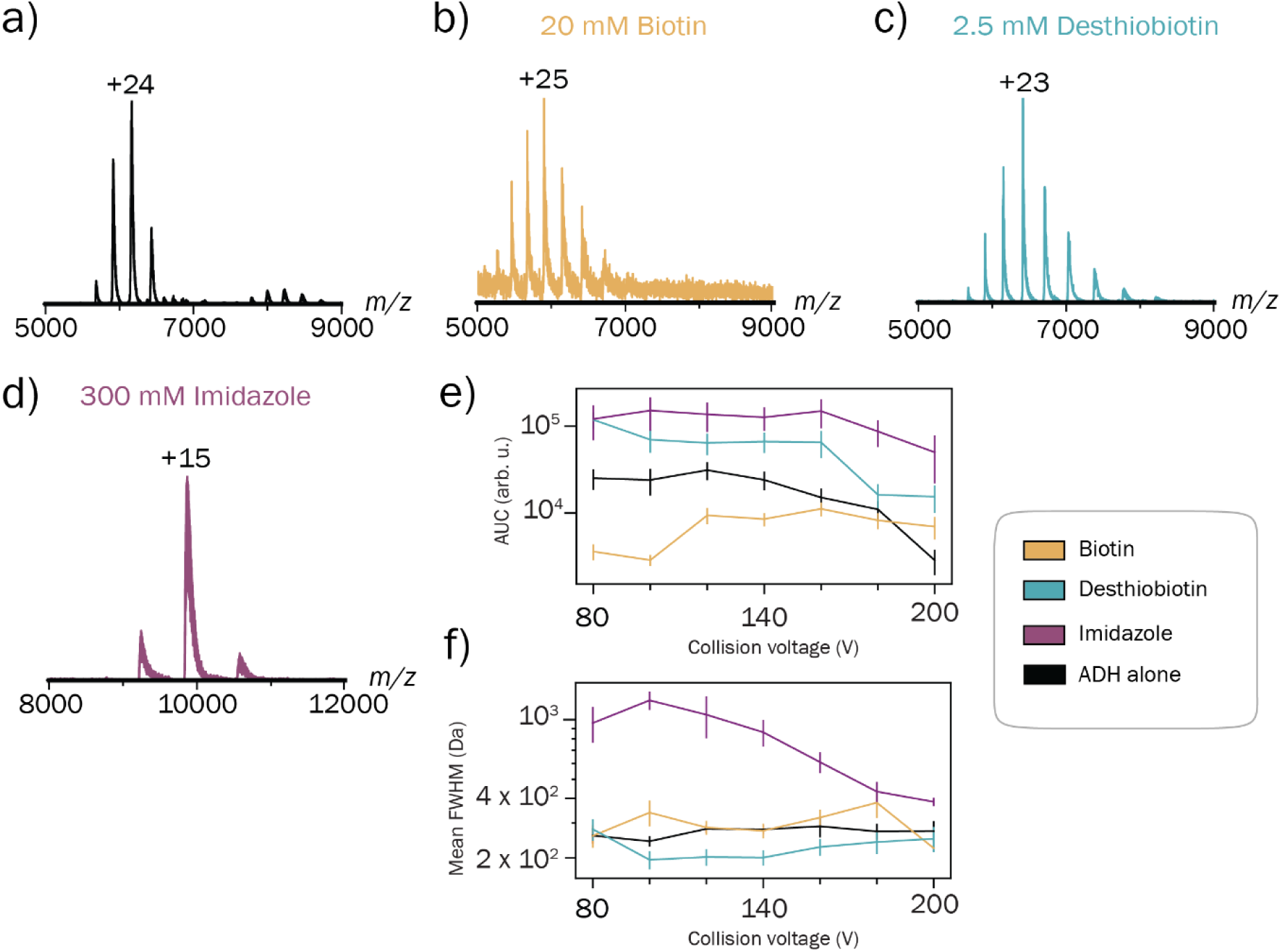
Spectra of the tetrameric ADH complex (148 kDa, 7.5 µM tetramer) in 150 mM AmAc acetate at pH 8 spiked with typical amounts of eluent at minimal collision voltages providing baseline resolved charge states on the Q-TOF 2. a) ADH at 5 V. b) ADH with 20 mM biotin at 100 V. c) ADH with 2.5 mM desthiobiotin at 80 V. d) ADH with 300 mM imidazole at 100 V. e) AUC and f) mean FWHM in regard to the CV at the respective eluent concentration.

**Figure 2:**
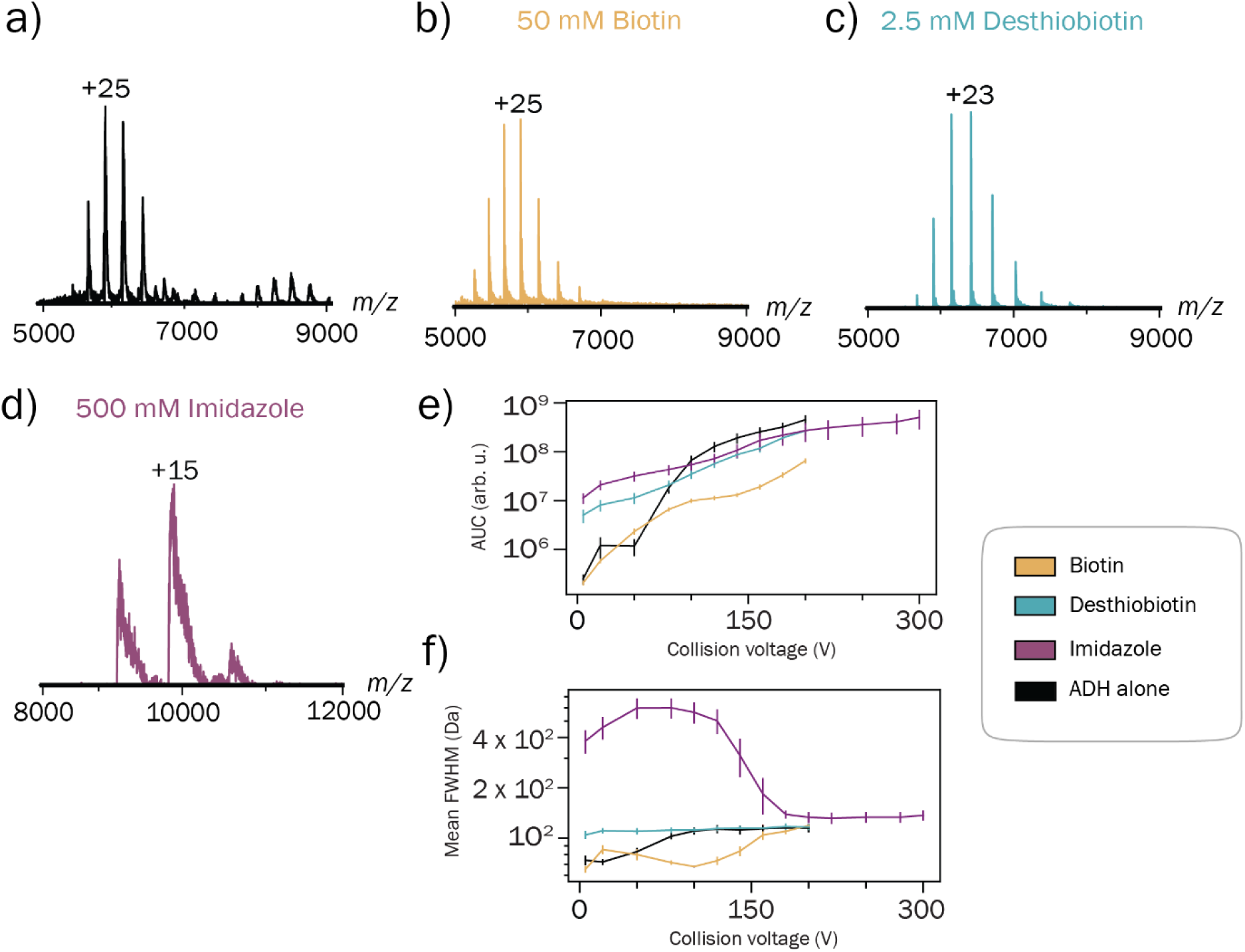
Spectra of the tetrameric ADH complex (148 kDa, 2.5 µM tetramer) in 150 mM AmAc at pH 8 spiked with typical amounts of eluent at minimal collision voltages providing baseline resolved charge states on the QExactive UHMR. a) ADH at 5 eV. b) ADH with 50 mM biotin at 140 eV. c) ADH with 2.5 mM desthiobiotin at 100 eV. d) ADH with 500 mM imidazole at 50 eV. e) AUC and f) mean FWHM in regard to the CV at the respective eluent concentration.

The graphs in Figure 1 and 2 e and f show the relation between area under the curve (AUC) and the mean FWHM of the charge states for each eluent. On the Q-TOF 2, the AUCs roughly stay constant (or show a slight increase for biotin) until around 160 V, when the signal broke down due to tetramer dissociation. However, the FWHM reduced significantly, which was in line with improved declustering, resulting in peak sharpening at elevated CV. On the QExactive UHMR, the situation was different. Due to nicely resolved peaks at low CV, the FWHM remained largely unaltered for biotin and desthiobiotin, but the transmission improved, resulting in an initial increase of the AUCs. For imidazole, we again observed peak sharpening, which was accompanied by an increase in AUC. This was probably due to the signal preprocessing in the QExactive UHMR, which is known to affect peak shape. Ultimately, both instruments produced high-quality spectra with high amounts of eluents, but the QExactive UHMR expectedly had an advantage at low protein amounts.

### Fast-track purification from different expression systems

#### Bacterial expression: Preparation and processing of viral polyprotein nsp7-11

The nsp7-11 polyprotein from SARS-CoV-2 contains regulatory factors of the enzymes driving viral replication and transcription, and its processing by the viral main protease (M^pro^) has previously been described.^30^ We chose this protein construct for proof-of-principle as it has been characterized and is available, both as His- and Strep-tagged versions, thus facilitating comparison of the affinity purification approaches. Moreover, it allowed us to test the compatibility of the workflow with enzymatic activity, subsequent oligomerization and retaining natural ligands like Zn^2+^ in nsp10.^38^ The polyprotein nsp7-11-His_6_ has a mass of 61.1 kDa, Strep-nsp7-11 has a mass of 61.8 kDa, both with a charge state distribution around 4000-5000 *m*/*z*.^30^ Spectra of the polyprotein were obtained on the QExactive UHMR (Figure 3 a-c and e-g) and the Q-TOF 2 (Figure 3 d and h). Measurements on the Q-TOF 2 were only performed for desthiobiotin, where the preferred elution concentration can be reached. The spectra in the presence of eluents were comparable to published data (UHMR,^30^ Q-TOF 2^39^). The spectra for both the polyprotein (left) and the processed sample (right) showed sharp peaks with good signal-to-noise. We found that samples with imidazole and desthiobiotin resulted in almost identical spectra compared to previous experiments with buffer exchange (Figure 3 a, e). We were also able to detect the nsp7+8 heterodimer and heterotetramer (Figure 3 b-h). Importantly, the nsp7+8 complexes have a particularly limited shelf-life and prolonged purification workflows often impair its formation. By avoiding buffer exchange and minimizing handling time, the fast-track workflow preserves the native state of the polyprotein and supports efficient processing and complex formation directly after elution, which is essential for reliably detecting the nsp7+8 heterodimer and heterotetramer. Moreover, nsp10 mass was in line with Zn^2+^ ligands being retained. In summary, the processing of the polyprotein with M^pro^ and the complexation of the resulting proteins were successful in the AmAc-based buffer surrogate with eluents present.

**Figure 3:**
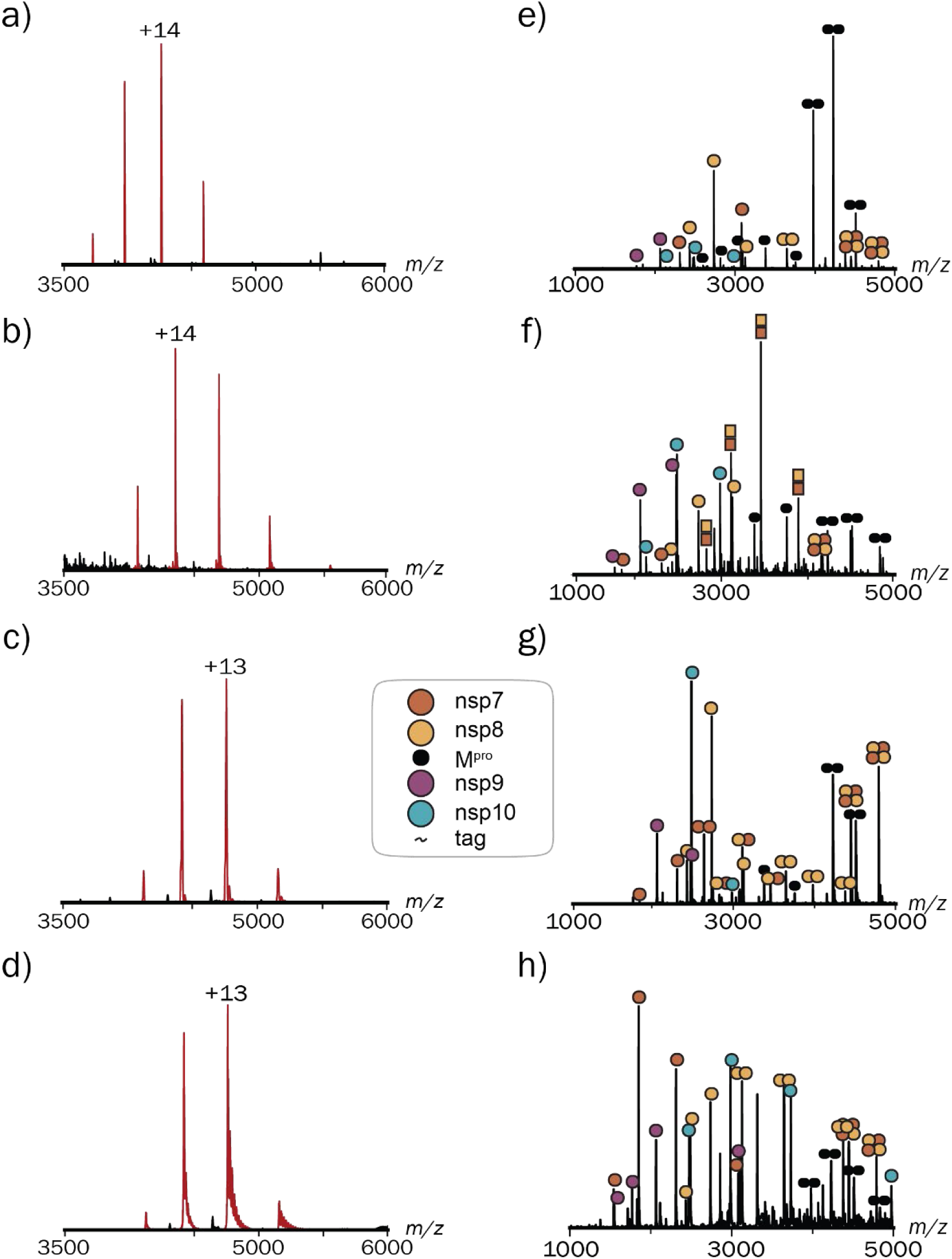
Spectra of 20 µM unprocessed (a-d) and different concentration of processed (e-h) nsp7-11 with M^pro^ without any eluent, and with imidazole or desthiobiotin after protein purification. a) nsp7-11-His_6_ without tag at 15 eV on the QExactive UHMR. b) nsp7-11-His_6_ with 100 mM imidazole at 15 eV on the QExactive UHMR. c) Strep-nsp7-11 with 1.5 mM desthiobiotin at 15 eV on the QExactive UHMR. d) Strep-nsp7-11 with 2 mM desthiobiotin at 50 V on the Q-TOF 2. e) processed nsp7-11 at 15 eV on the QExactive UHMR. f) processed nsp7-11-His_6_ with 75 mM imidazole at 15 eV on the QExactive UHMR. g) 13 µM Strep-nsp7-11 with 3 µM M^pro^ and 0.75 mM desthiobiotin at 10 eV on QExactive UHMR. h) 15 µM Strep-nsp7-11 with 3 µM M^pro^ and 1 mM desthiobiotin at 100 V on the Q-TOF 2.

#### Comparing eukaryotic and bacterial expression: Preparation of the viral polymerase nsp12

SARS-CoV-2 nsp12 is the central RNA-dependent RNA polymerase (RdRp) of the viral replication-transcription complex, synthesizing genomic and subgenomic RNA together with cofactors nsp7 and nsp8 and further nsps.^40^ To assess the fast-track workflow on a large multidomain enzyme, we used nsp12 as a test case. We compared material purified using the conventional multi-step protocol described by Chen *et al*. (2020)^41^, which involves Heparin, His-tag, size-exclusion chromatography, and an additional buffer exchange step with nsp12 obtained directly through the streamlined fast-track procedure using Strep-tag affinity chromatography. Using the traditional purification and buffer-exchange workflow, 10 µg nsp12 were recovered from 1 L *E. coli* culture. The resulting spectrum shows clear charge states contained elevated background signals and additional peaks from co-purifying bacterial proteins (Figure 4a). Moreover, the expression, purification and buffer exchange were hardly reproducible. Only occasionally sufficient yields for subsequent analysis could be obtained. In contrast, the fast-track approach reproducibly gave good yields with substantially increased recovery with 717 µg nsp12 from 500 mL *E. coli* culture and 114 µg from mammalian expression (Figure 4b, c). The measured masses further support the assigned nsp12 species. The conventionally purified His-SUMO-nsp12 corresponds to the tag-cleaved protein carrying a DTT-derived adduct, whereas the HEK293-derived Twin-Strep-nsp12 is consistent with an Fe³⁺-bound species (Table S6).^42,43^ Both preparations produced highly resolved spectra with reduced chemical background, demonstrating that the minimal handling preserves the native structural integrity of the polymerase. As observed for nsp7-11, the presence of desthiobiotin did not affect the stability or spectral properties of nsp12. Overall, the fast-track workflow enables efficient purification of nsp12 across expression systems, provides markedly higher yield, and delivers high-quality native mass spectra directly after elution.

**Figure 4:**
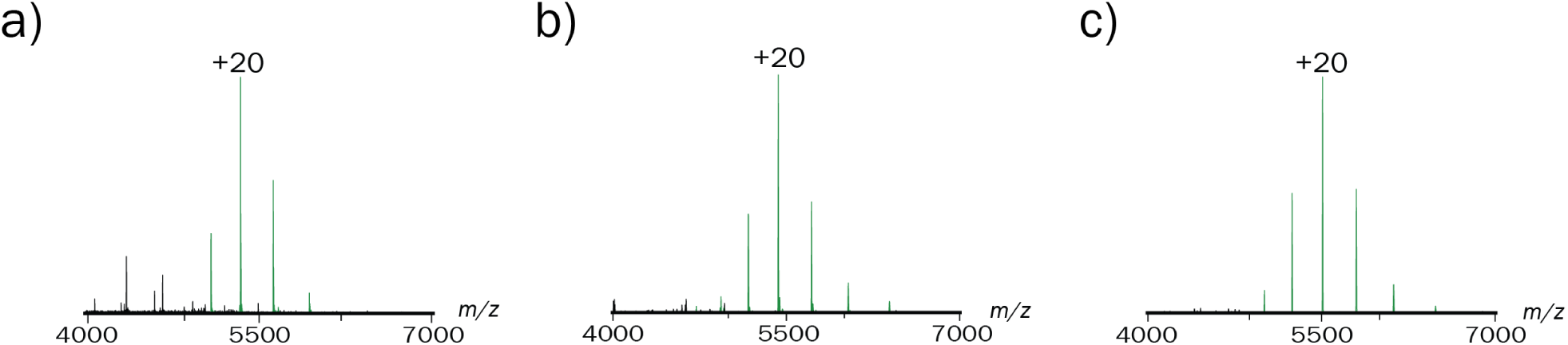
Spectra of nsp12 purified via fast-track protocol a) bacterially expressed His_6_-SUMO-nsp12 after enzymatic digest via ULp1 in 300 mM AmAc 2 mM DTT (5 µM) purified via multistep approach at 50 eV (with no technical replica, b) bacterially expressed Strep-nsp12 (10 µM) with 2.5 mM desthiobiotin at 80 eV, c) eukaryotically expressed of nsp12-Twin-Strep (1 µM) with 2.5 mM desthiobiotin at 0 eV.

#### Eukaryotic expression: Preparation of the viral N-protein and analysis of prost-translational modifications

The SARS-CoV-2 nucleocapsid (N) protein plays a central role in RNA genome packaging and regulation of replication, and is subject to extensive phosphorylation that modulates its interactions and oligomerization behavior.^44^ To further evaluate the performance of the fast-track workflow for mammalian expression, we directly compared N-protein prepared using conventional affinity-purification followed by buffer exchange with material obtained through the streamlined fast-track procedure. Using the classical workflow, 43 µg N-protein were recovered after the buffer exchange, yielding spectra with sharp peaks and low chemical background (Figure 5 a). The fast-track preparation produced considerably more protein (up to 200 µg in one fraction) but also retained additional co-eluting impurities. Importantly, the fast-track spectra revealed a more complete set of N-protein species, including an additional phosphorylated isoform that was not detected after the conventional purification (Figure 5 b). Despite the higher background, the fast-track workflow therefore provides superior overall performance, primarily due to its markedly higher yield and its ability to preserve the natural heterogeneity of post-translationally modified N-protein. Together, these findings show that the fast-track workflow substantially improves yield and enhances the detectability of post-translationally modified N-protein species, even when accompanied by minor additional background signals.

**Figure 5:**
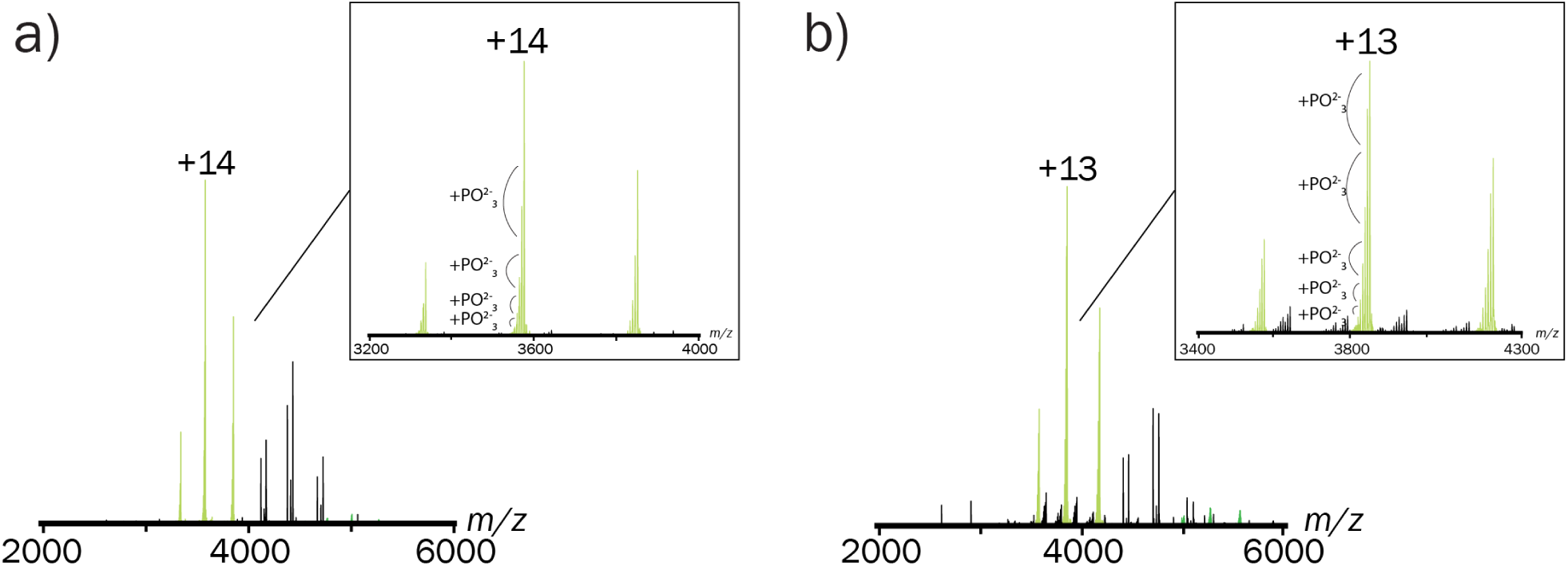
Spectra of the N-protein (50 kDa, 8 µM) in 500 mM AmAc at pH 8 a) using conventional Tris-buffers with buffer exchange step, b) using fast-track purification method at 100 eV collision voltages on the QExactive UHMR.

#### Eukaryotic expression systems: Stoichiometry of herpes viral proteins

Next, the protocol was tested with viral protein complexes from herpes simplex virus 1 (HSV-1) and KSHV (Kaposi’s sarcoma-associated herpesvirus) from eukaryotic expression systems. We determined the stoichiometries of the viral packaging motor (terminase) of HSV-1, its associated protein pUL32 (HSV-1) as well as its homolog pORF68 from KSHV. The terminase is a heterotrimer consisting of pUL15, pUL28, and pUL33 (HSV-1, strain 17). Protein yields from mammalian and insect cells are usually lower than in *E. coli*. This can be troublesome when analyzing large protein complexes by nMS, which require sufficiently high protein concentrations for higher-order complex formation. To purify the terminase a His-pUL15 construct was used, nevertheless protein yields from insect cells after conventional sample preparation were insufficient for nMS. By using the fast-track, omitting SEC and buffer exchange, we obtained a fairly high protein yield of 3 mg for the mixture of all three proteins in AmAc supplemented with 350 mM imidazole. This allowed for ten-fold dilution, eliminating imidazole impact on the native mass spectra. As shown in Figure 6 a, we found monomers, dimers, and trimers of the HSV-1 terminase heterotrimer. The remaining mass discrepancies are likely related to the complex and still incompletely characterized nature of the terminase, for which PTMs, ligand binding, or reported truncations of the main subunit may contribute to the observed masses (Table S8). The broad peaks and increased FWHM values further support this interpretation (Figure 6, Table S8). Sequence drift of the constructs is unlikely, as all plasmids were sequence-verified after purchase and protein products remained unchanged over time based on SDS-PAGE analysis (Figure S7). Next, we measured a mix of separately purified HSV-1 terminase with Strep-UL32 both derived from insect cells (Figure 6 b). We observed again monomeric terminase heterotrimers as well as homotrimers of Strep-UL32. Currently, it remains unclear if this UL32 homotrimer is a physiological form or an artifact of the expression in insect cells. Electron microscopy data suggest pentamers of UL32 from mammalian expression.^45^ Finally, TwinStrep-ORF68 expressed in mammalian cells resulted in decamers, likely dimers of pentamers (Figure 6c) in line with previous electron microscopy data.^45^ In conclusion, we showed that our fast-track approach is ideal to obtain enough material from insect and mammalian cells for nMS, ensuring correct protein folding and potentially important and physiological PTMs.

**Figure 6:**
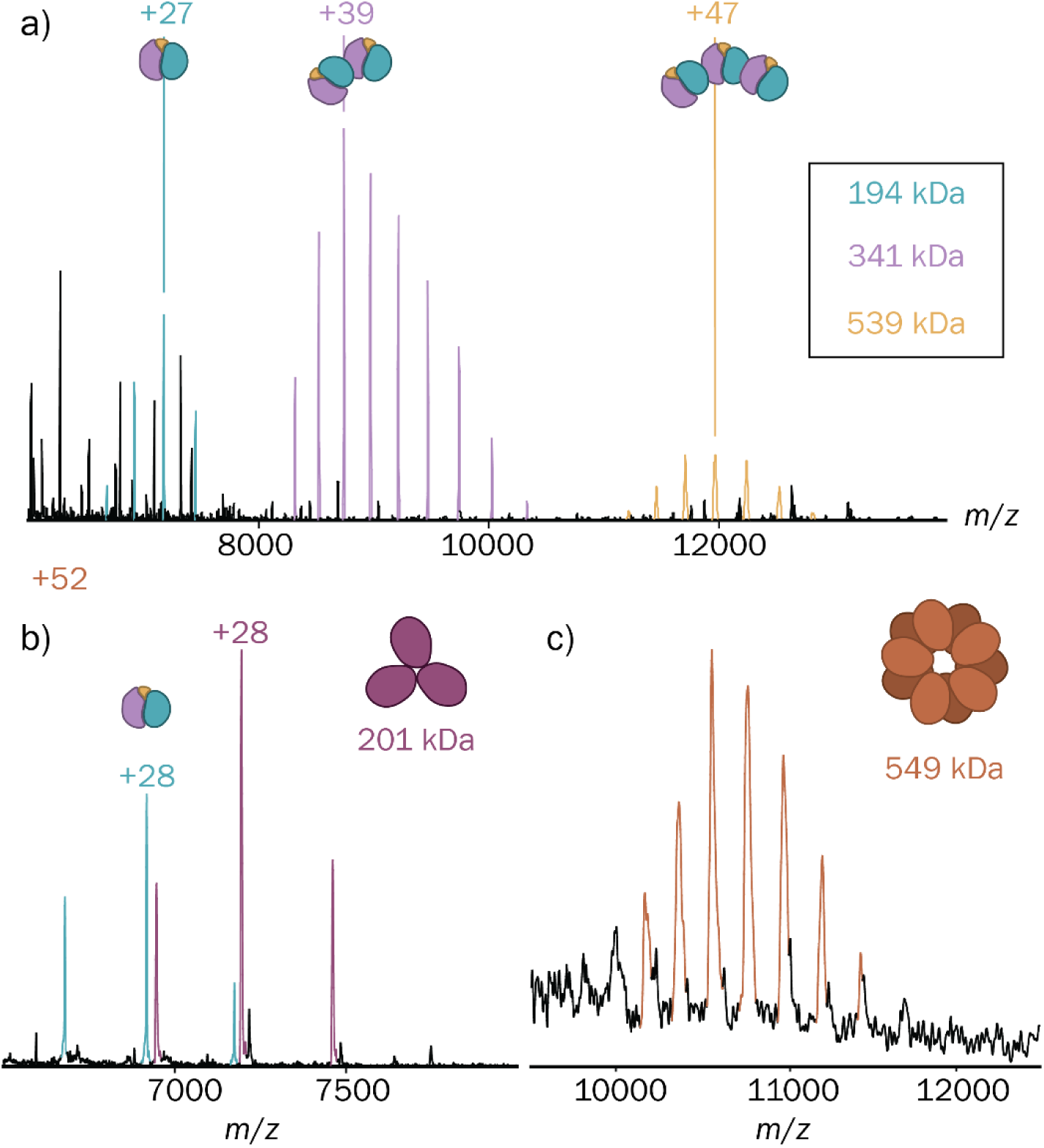
Representative spectra of different herpes virus proteins expressed in insect or mammalian cells. Proteins were purified via the fast-track protocol and measured on a QExactive UHMR. a) 3 µM HSV-1 terminase heterotrimer expressed in insect cells with 35 mM imidazole purified via His-tag with higher-energy collisional dissociation (HCD) turned off to avoid disrupting complex formation. b) 5 µM Strep-pUL32 (HSV-1) expressed in insect cells mixed with 1 µM terminase heterotrimer, and 2.5 mM desthiobiotin at 250 eV. c) 5 µM pORF68 (KSHV) expressed in mammalian cells and purified via Twin-Strep with 2.5 mM desthiobiotin at 250 eV (all having no technical replica).

## Conclusion

Using the described fast-track sample preparation protocol, we are able to obtain high-quality nMS spectra for proteins from different expression systems. After lysis, nMS was obtained within 2 h where common protein purification protocols for nMS analysis take 1.5-2 days. This included an affinity purification step, which is usually necessary for low-abundant or endogenously expressed proteins. This also holds if the protein is tagged or produced in infection context. The shortened sample preparation time is particularly advantageous for flexible proteins with low stability, which often possess limited shelf-lives. Besides determining the stoichiometry of the protein complexes, we also were able to monitor enzymatic activity and protein complexation in an nMS-suitable buffer surrogate. Three eluents (imidazole, biotin, desthiobiotin) that are commonly used in standard protein purification were tested and all were compatible. In general, the QExactive UHMR setup performed better at high eluent concentrations, but spectra obtained from lower concentrations were qualitatively indistinguishable from data obtained on a TOF system. Superior performance was observed with Strep-tag/desthiobiotin system due to higher specificity of the tag and high resolution at maximum required eluent concentration without limiting collisional activation range for MS/MS.

For example, Ni-based resins are known to copurify proteins such as transferrin from expression media of mammalian expression systems.^12,46^ In conjunction with the low target protein yields, the high ionization propensity and mass heterogeneity as a glycoprotein hamper target analysis. Besides, imidazole is also a charge reducing agent that can help with protein stabilization during ionization but reduces the stability of proteins when kept in the storage buffer.^35^ Hence, while one of the aims was a fast purification strategy, imidazole-based elution makes fast analysis a necessity. Imidazole also limited the remaining activation range for MS/MS of protein complexes due to enhanced required desolvation. Protein purification from *E. coli* often results in high yields compatible with His-tag based purification and multi-fold dilutions, resulting in charge state resolved spectra without excessive activation and independent of the employed mass spectrometer. To conclude, we recommend the use of Strep tags for protein purification with desthiobiotin as eluent for fast-track purification for in-depth nMS analysis of low yield or low stability expressing proteins from eukaryotic expression systems. Moreover, we suspect that other affinity purification strategies based on small eluents such as glutathione and flag would be compatible with the approach.

## Supporting information

Complete supplemental information

## Acknowledgements

AFRG, JB, and CU acknowledge funding from the Landesforschungsförderung Hamburg (LFF) through the ZeMeIn project. FAS acknowledges funding from Avicenna-Studienwerk. CU acknowledges funding from the Leibniz Association through SAW-2014-HPI-4 grant and the BMFTR VirMScan 13GW0622. CU, KSK and AFRG acknowledge funding through H2020 ERC-StG SPOCk’S MS No. 759661. JB is funded by the Deutsche Forschungsgemeinschaft (DFG, German Research Foundation) under Germany’s Excellence Strategy - EXC 2155 - project number 390874280. A.B. and C.U. acknowledge SPIDoc’s – The next generation MS SPIDoc’s – Marie Skłodowska Curie Action program HORIZON-MSCA-2022-DN. J.B. and C.U. acknowledge funding by the DFG-funded RTG 2771 Humans and Microbes, project number 453548970, the DFG-funded RTG 2887 VISION, project number 49735088, DFG-funded CRC 1648 Emerging Viruses project number SFB 1648/1 2024-512741711, the Leibniz ScienceCampus InterACt, funded by the BWFGB Hamburg and the Leibniz Association (W75/2022) InterACt and “Hamburg-X Infektionsforschung”. We thank Britt Glaunsinger for the kind gift of pcDNA4/TO-2xStrep-ORF68 through Addgene (Addgene plasmid #162625; http://n2t.net/addgene:162625; RRID: Addgene_162625)). The Leibniz Institute of Virology (LIV) is supported by the Free and Hanseatic City of Hamburg and the Federal Ministry of Health.

