## Supplementary material for "Fast tracking native mass spectrometry: Skipping over buffer exchange": Complete supplemental information

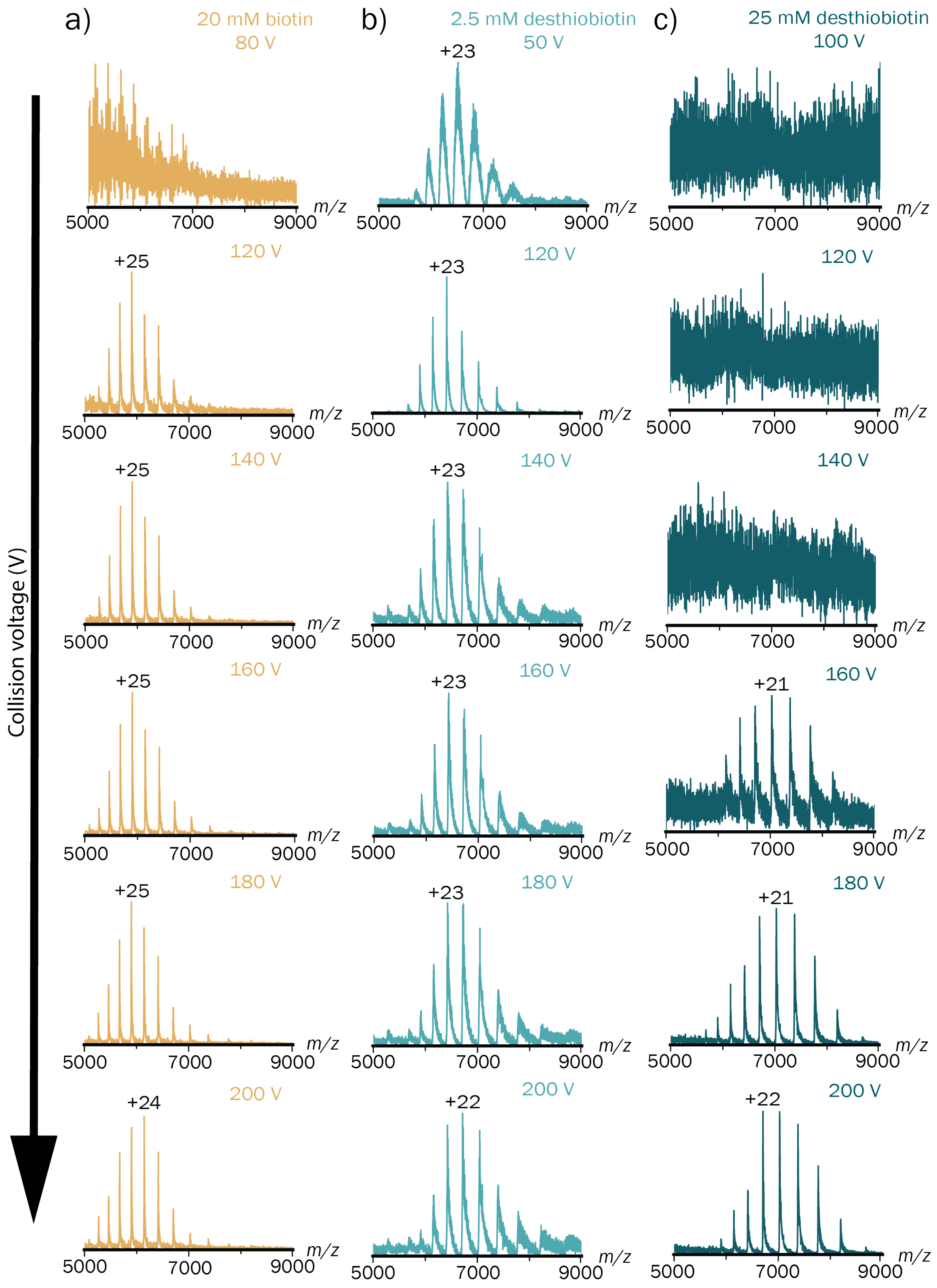


*Figure S1:* Spectra of the tetrameric ADH complex (148 kDa, 2.5 µM) in 150 mM ammonium acetate at pH 8 spiked with typical amounts of eluent at different collision voltages, increasing from top to bottom, on the Q-ToF2. a) ADH and 20 mM biotin at 80-200 V. b) ADH and 2.5 mM desthiobiotin at 50-200 V. c) ADH and 25 mM desthiobiotin at 100-200 V

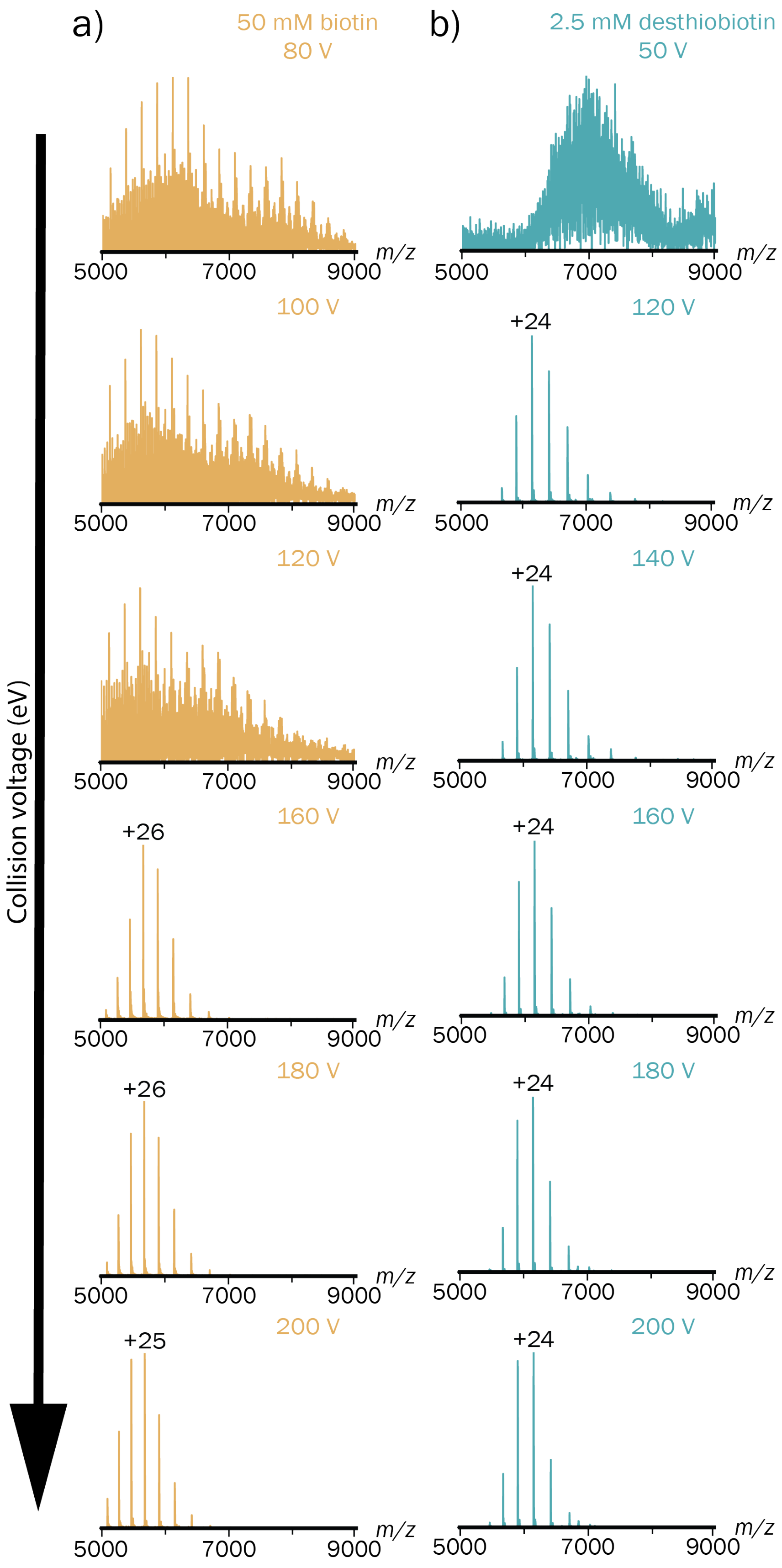


*Figure S2:* Spectra of the tetrameric ADH complex (148 kDa, 2.5 µM) in 150 mM ammonium acetate at pH 8 spiked with typical amounts of eluent at different collision voltages, increasing from top to bottom, on the QExactive UHMR 2. a) ADH and 50 mM biotin at 80-200 V. b) ADH and 2.5 mM desthiobiotin at 50-200 V.

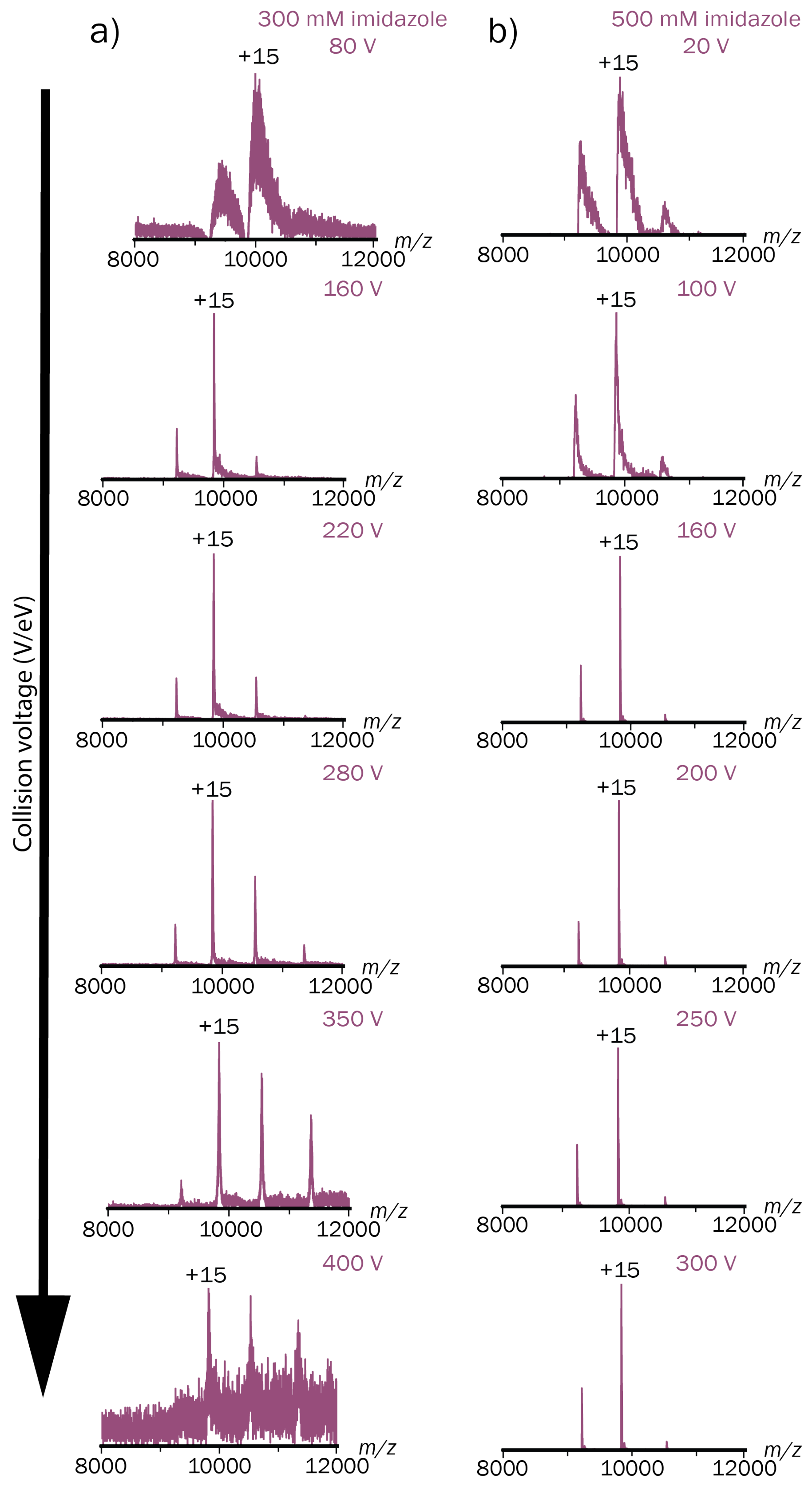


*Figure S3:* Spectra of the tetrameric ADH complex (148 kDa, 2.5 µM) in 150 mM ammonium acetate at pH 8 spiked with typical amounts of the eluent imidazole at different collision voltages, increasing from top to bottom, analyzed on two different mass spectrometers. **a)** ADH and 300 mM imidazole at 80-400 V on the Q-ToF2. **b)** ADH and 500 mM imidazole at 20-300 V on the QExactive UHMR.

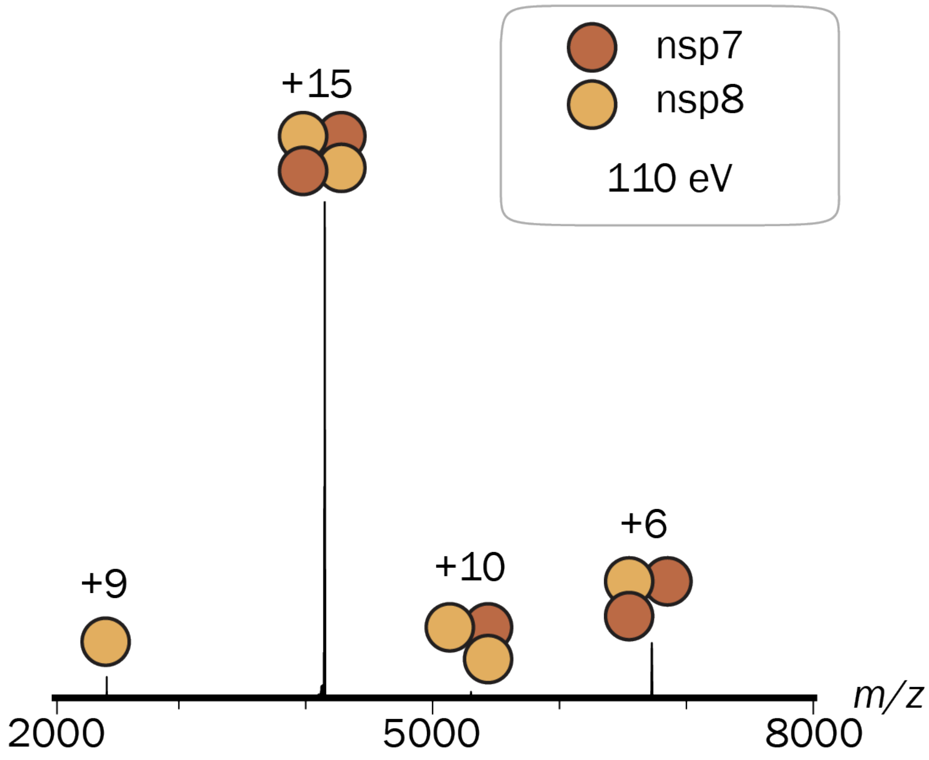


*Figure S4:* MS2 spectrum of the heterotetrameric nsp7+nsp8 complex (62 kDa, 15 µM) of the nsp7-11-His purification in 300 mM ammonium acetate with 1 mM dithiothreitol at pH 8 in the presence of 3 µM Mpro and 90 mM imidazole at 110 eV measured on the QExactive UHMR.

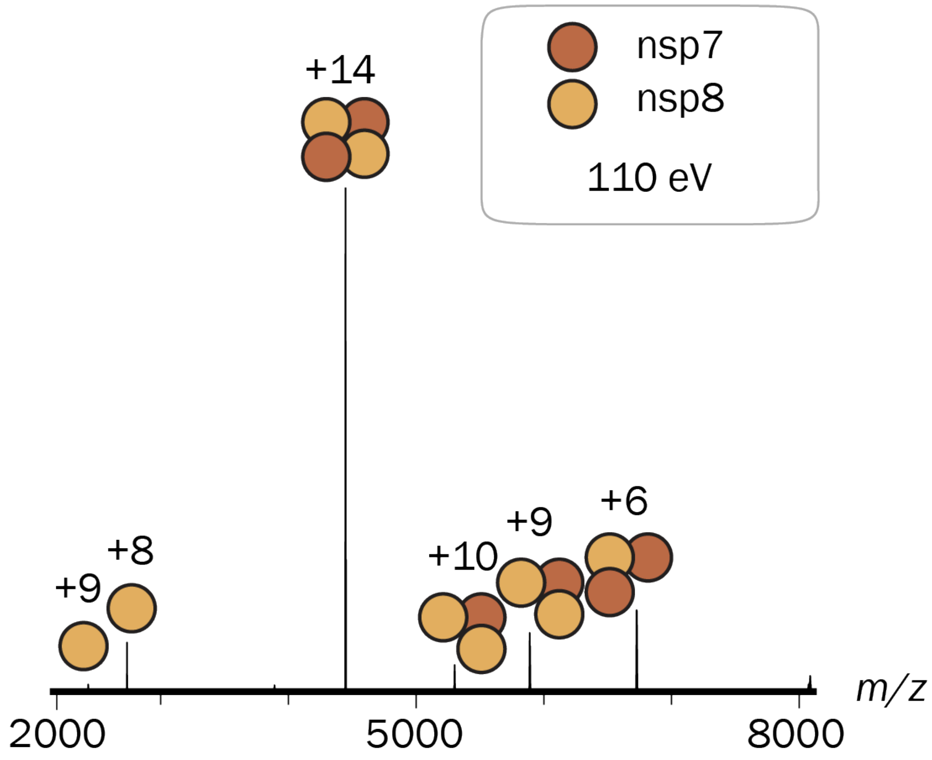


*Figure S5:* MS2 spectrum of the heterotetrameric nsp7+nsp8 complex (62 kDa, 12.6 µM) of the Strep-nsp7-11 purification in 300 mM ammonium acetate with 1 mM dithiothreitol at pH 8 in the presence of 3.5 µM Mpro and 1.25 mM desthiobiotin at 110 eV measured on the QExactive UHMR.

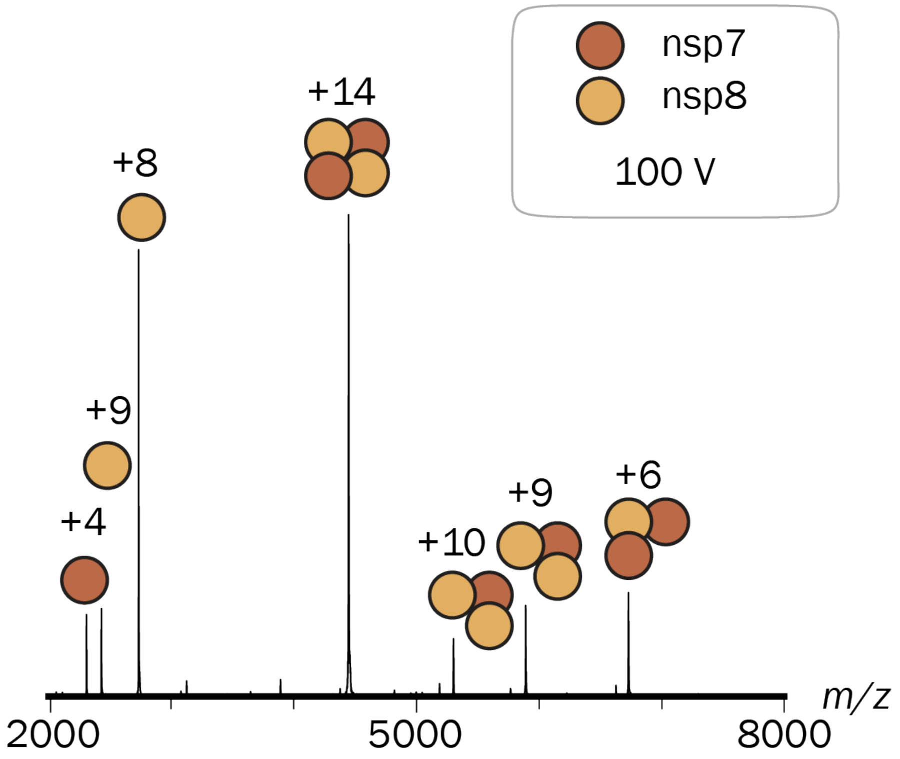


*Figure S6:* MS2 spectrum of the heterotetrameric nsp7+nsp8 complex (62 kDa, 15 µM) of the Strep-nsp7-11 purification in 300 mM ammonium acetate with 1 mM dithiothreitol at pH 8 in the presence of 3 µM Mpro and 1.25 mM desthiobiotin at 100 V measured on the Q-ToF2.

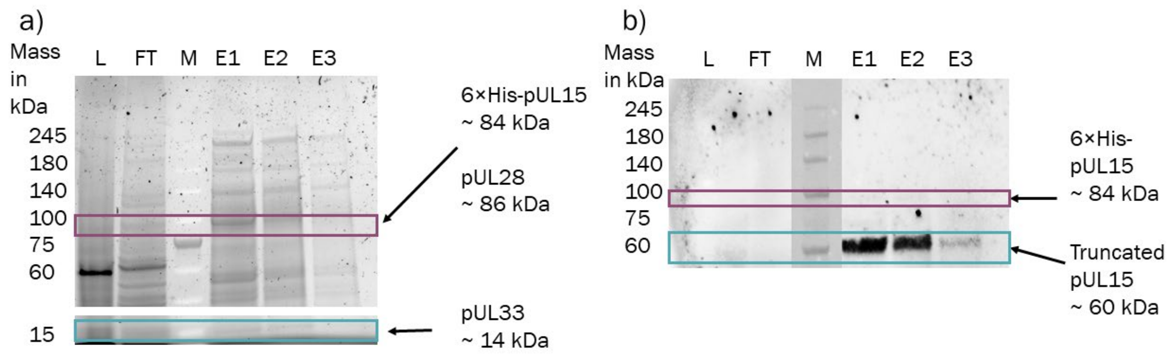


*Figure S7:* a) SDS-Page and b) western-blot of Fractions of the fast-track purification of the terminase with L being the lysate, FT being the flow-through, M being the marker molecular weights in kDa and E1-3 being the elutions in their respective order.

*Table S1:* ADH tetrameric complex ($ADH_4$) with and without elution substances were detected on the Q-ToF2. FWHM and measured masses were averaged and standard error is given.[1]

| Protein | Th. Mass (in Da) | Measured Mass (in Da) | FWHM (in Da) |
|---|---|---|---|
| $ADH_4$ alone | 146824 | 147930 ± 180 | 770 ± 220 |
| $ADH_4$+desthiobiotin (2.5mM) | 146824 | 148000 ± 500 | 1300 ± 700 |
| $ADH_4$+imidazole (300mM) | 146824 | 147970 ± 270 | 1440 ± 270 |
| $ADH_4$+biotin (20mM) | 146824 | 147650 ± 70 | 720 ± 130 |

*Table S2:* ADH tetrameric complex ($ADH_4$) with and without elution substances were detected on the Q Exactive UHMR Orbitrap. FWHM and measured masses were averaged and standard error is given.[1]

| Protein | Th. Mass (in Da) | Measured Mass (in Da) | FWHM (in Da) |
|---|---|---|---|
| $ADH_4$ alone | 146824 | 147730 ± 170 | 140 ± 140 |
| $ADH_4$+desthiobiotin (2.5mM) | 146824 | 147550 ± 20 | 130 ± 10 |
| $ADH_4$+imidazole (300mM) | 146824 | 148300 ± 400 | 1700 ± 400 |
| $ADH_4$+biotin (20mM) | 146824 | 147560 ± 10 | 101 ± 4 |

*Table S3:* Strep-nsp7-11 polyprotein and all mass species were detected on the Q-ToF2. FWHM and measured masses were averaged and standard error is given.[2]

| Protein | Th. Mass (in Da) | Measured Mass (in Da) | FWHM (in Da) |
|---|---|---|---|
| StrepII-nsp7-11 polyprotein+2$Zn^{2+}$ | 61874 | 61881 ± 1 | 29 ± 7 |
| $M^{pro}$-His dimer | 67337<br>67645.5<br>(+ DTT adducts) | 67611 ± 1 | 34 ±12 |
| nsp7 | 9240 | 9244.1 ± 0.2 | 7.2 ± 0.3 |
| nsp8 | 21881 | 21889.7 ± 0.4 | 11.4 ± 0.3 |
| nsp8-dimer | 43762 | 43773.7 ± 0.6 | 23 ± 3 |
| nsp9 | 12378 | 12384.6 ± 0.1 | 8.7 ± 0.2 |
| nsp10+2$Zn^{2+}$ | 14920 | 14922.0 ± 0.2 | 10.3 ± 0.4 |
| nsp7+8 heterotetramer | 62242 | 62256.4 ± 0.7 | 28 ± 5 |

*Table S4:* Strep-nsp7-11 polyprotein and all mass species were detected on the QExactive UHMR Orbitrap. FWHM and measured masses were averaged and standard error is given.[2]

| Protein | Th. Mass (in Da) | Measured Mass (in Da) | FWHM (in Da) |
|---|---|---|---|
| StrepII-nsp7-11 polyprotein+2$Zn^{2+}$ | 61874 | 61889 ± n.d. | 43 ± n.d. |
| $M^{pro}$-His dimer | 67337<br>67645.5<br>(+ 2 DTT adducts) | 67614 ± 2 | 37 ± 2 |
| nsp7 | 9240 | 9243.23 ± 0.08 | 6.57 ± 0.05 |
| nsp8 | 21881 | 21889.2 ± 0.3 | 10.6 ± 0.3 |

| | | | |
|---|---|---|---|
| nsp8 dimer | 43762 | 43773.6 ± 1.3 | 24.1 ± 1.7 |
| nsp9 | 12378 | 12383.3 ± 0.2 | 7.4 ± 0.2 |
| nsp10+ $2Zn^{++}$ | 14920 | 14921.51 ± 0.12 | 9.90 ± 0.02 |
| nsp7+8 heterotetramer | 62242 | 62260.3 ± 1.0 | 36.7 ± 1.3 |

*Table S5:* nsp7-11-His polyprotein and all mass species were detected on the QExactive UHMR Orbitrap. FWHM and measured masses were averaged and standard error is given.[2]

| Protein | Th. Mass (in Da) | Measured Mass (in Da) | FWHM (in Da) |
|---|---|---|---|
| nsp7-11-His polyprotein+$2Zn^{+2}$ | 60954 | 60982 ± n.d. | 45 ± n.d. |
| $M^{pro}$ | 33669<br>33823.3<br>(+DTT adduct) | 33805.5 ± 0.8 | 15.5 ± 0.1 |
| $M^{pro}$ dimer | 67337<br>67645.5<br>(+ 2 DTT adducts) | 67648 ± 25 | 58 ± 20 |
| nsp7 | 9240 | 9243.7± 0.1 | 8.2 ± 0.4 |
| nsp8 | 21881 | 21889.1 ± 0.3 | 15 ± 3 |
| nsp9 | 12378 | 12382.9 ± 0.1 | 9.0 ± 0.1 |
| nsp10+$Zn_2$ | 14920 | 14920 ± 4 | 9.5 ± 0.3 |
| nsp7+8 heterotetramer | 62242 | 62268 ± 5 | 62 ± 26 |
| nsp7-8 polyprotein | 31103 | 31112.5 ± 0.2 | 16.8 ± 0.6 |

*Table S6:* nsp12 was detected on the QExactive UHMR Orbitrap. FWHM and measured masses were averaged and standard error is given.[3–5]

| Protein | Th. Mass (in Da) | Measured Mass (in Da) | FWHM (in Da) |
|---|---|---|---|
| His-SUMO-nsp12 (*E. coli*) | 120357 (with tag)<br>106660 (without tag)<br>16812 (+DTT adduct) | 106809 ± n.d. | 36 ± n.d. |
| Strep-nsp12 (*E. coli*) | 108602 | 108619 ± 4 | 71 ± 8 |
| Twin-Strep-nsp12 (HEK293) | 110117<br>110170 (+$Fe^{3+}$) | 110170 ± 3 | 74 ± 9 |

*Table S7:* N-protein was detected on the QExactive UHMR Orbitrap. FWHM and measured masses were averaged and standard error is given.[6]

| Protein | Th. Mass (in Da) | Measured Mass (in Da) | FWHM (in Da) |
|---|---|---|---|
| N-protein (fast-track protocol) | 48951<br>(without PTMS)<br>50071<br>(+14 phosphorylation) | 50074.5 ± 0.6 | 19 ± 3 |
| N-protein (regular protocol) | 48951<br>(without PTMS)<br>50071<br>(+14 phosphorylation) | 50075.0 ± 0.5 | 16.8 ± 0.7 |

*Table S8:* The protein complexes of pORF68 (KSHV), Strep-pUL32 (HSV-1) and the terminase (HSV-1) were detected on the QExactive UHMR Orbitrap. FWHM and measured masses were averaged and standard error is given. Triplicates were recorded for pUL32 and the terminase complex. For pORF68 only a singular spectra was recorded.

| Protein | Th. Mass (in Da) | Measured Mass (in Da) | FWHM (in Da) |
|---|---|---|---|
| terminase complex | 183973 (without PTMS) | 193774 ± 1 | 189 ± 7 |
| dimer of terminase complex | 367946 (without PTMS) | 340968 ± 1 | 280 ± 18 |
| trimer of terminase complex | 551919 (without PTMS) | 538600 ± 100 | 1060 ± 110 |
| Trimer of pUL32 | 201183 (without PTMS) | 201435 ± 14 | 129 ± 25 |
| pORF68 decamer complex | 550970 (without PTMS) | 549180 ± 30 | 1700 ± 500 |

*Table S9:* Amino acid sequences and theoretical masses of recombinantly expressed proteins.

| Protein | Sequence | Theoretical Mass (Da) |
|---|---|---|
| nsp7-11-His | SKMSDVKCTSVVLLSVLQQLRVESSSKLWAQCVQLHNDIL LAKDTTEAFEKMVSLLSVLLSMQGAVDINKLCEEMLDNR ATLQAIASEFSSLPSYAAFATAQEAYEQAVANGDSEVVLK KLKKSLNVAKSEFDRDAAMQRKLEKMADQAMTQMYKQ ARSEDKRAKVTSAMQTMLFTMLRKLDNDALNNIINNA RDGCVPLNIIPLTTAAKLMVVIPDYNTYKNTCDGTTFTYAS ALWEIQQVVDADSKIVQLSEISMDNSPNLAWPLIVTALRAN SAVKLQNNELSPVALRQMSCAAGTTQTACTDDNALAYYN TTKGGRFVLALLSDLQDLKWARFPKSDGTGTIYTELEPPCR FVTDTPKGPKVKYLYFIKGLNNLNRGMVLGSLAA TVRLQAGNATEVPANSTVLSFCAFAVDAAKAYKDYLASG GQPITNCVKMLCTHTGTGQAITVTPEANMDQESFGGASCC LYCRCHIDHPNPKGFCDLKGKYVQIPTTCANDPVGFTLKN TVCTVCGMWKGYGCSCDQLREPMLQSADAQSFLNGFAVS ARGSHHHHHH | 60824 |
| Strep-nsp7-11 | MASWSHPQFEKGGSAVLQSKMSDVKCTSVVLLSVLQQLR VESSSKLWAQCVQLHNDILLAKDTTEAFEKMVSLLSVLLS MQGAVDINKLCEEMLDNRATLQAIASEFSSLPSYAAFATA QEAYEQAVANGDSEVVLKKLKKSLNVAKSEFDRDAAMQ RKLEKMADQAMTQMYKQARSEDKRAKVTSAMQTMLFT MLRKLDNDALNNIINNARDGCVPLNIIPLTTAAKLMVVIPD YNTYKNTCDGTTFTYASALWEIQQVVDADSKIVQLSEISM DNSPNLAWPLIVTALRANSAVKLQNNELSPVALRQMSCAA GTTQTACTDDNALAYYNTTKGGRFVLALLSDLQDLKWAR FPKSDGTGTIYTELEPPCRFVTDTPKGPKVKYLYFIKGLNN LNRGMVLGSLAATVRLQAGNATEVPANSTVLSFCAFAVD AAKAYKDYLASGGQPITNCVKMLCTHTGTGQAITVTPEAN MDQESFGGASCCLYCRCHIDHPNPKGFCDLKGKYVQIPTT CANDPVGFTLKNTVCTVCGMWKGYGCSCDQLREPMLQS ADAQSFLNGFAVSAET | 61874 |
| His-SUMO- | MGSSHHHHHHSSGLVPRGSHMASMSDSEVNQEAKPEVKP EVKPETHINLKVSDGSSEIFFKIKKTTPLRRLMEAFAKRQG | 120357 |

| | | |
|---|---|---|
| nsp12 | KEMDSLRFLYDGIRIQADQTPEDLDMEDNDIIEAHREQIGG<br>SADAQSFLNRVCGVSAARLTPCGTGTSTDVVYRAFDIYND<br>KVAGFAKFLKTNCCRFQEKDEDDNLIDSYFVVKRHTFSNY<br>QHEETIYNLLKDCPAVAKHDFFKFRIDGDMVPHISRQRLTK<br>YTMADLVYALRHFDEGNCDTLKEILVTYNCCDDDYFNKK<br>DWYDFVENPDILRVYANLGERVRQALLKTVQFCDAMRNA<br>GIVGVLTLDNQDLNGNWYDFGDFIQTTPGSGVPVVDSYYS<br>LLMPILTLTRALTAESHVDTDLTKPYIKWDLLKYDFTEERL<br>KLFDRYFKYWDQTYHPNCVNCLDDRCILHCANFNVLFST<br>VFPPTSFGPLVRKIFVDGVPFVVSTGYHFRELGVVHNQDV<br>NLHSSRLSFKELLVYAADPAMHAASGNLLLDKRTTCFSVA<br>ALTNNVAFQTVKPGNFNKDFYDFAVSKGFFKEGSSVELKH<br>FFFAQDGNAAISDYDYYRYNLPTMCDIRQLLFVVEVVDKY<br>FDCYDGGCINANQVIVNNLDKSAGFPFNKWGKARLYYDS<br>MSYEDQDALFAYTKRNVIPTITQMNLKYAISAKNRARTVA<br>GVSICSTMTNRQFHQKLLKSIAATRGATVVIGTSKFYGGW<br>HNMLKTVYSDVENPHLMGWDYPKCDRAMPNMLRIMASL<br>VLARKHTTCCSLSHRFYRLANECAQVLSEMVMCGGSLYV<br>KPGGTSSGDATTAYANSVFNICQAVTANVNALLSTDGNKI<br>ADKYVRNLQHRLYECLYRNRDVDTDFVNEFYAYLRKHFS<br>MMILSDDAVVCFNSTYASQGLVASIKNFKSVLYYQNNVF<br>MSEAKCWTETDLTKGPHEFCSQHTMLVKQGDDYVYLPYP<br>DPSRILGAGCFVDDIVKTDGTLMIERFVSLAIDAYPLTKHP<br>NQEYADVFHLYLQYIRKLHDELTGHMLDMYSVMLTNDNT<br>SRYWEPEFYEAMYTPHTVLQ | |
| Strep-<br>nsp12 | MASWSHPQFEKGGSAVLQSADAQSFLNRVCGVSAARLTP<br>CGTGTSTDVVYRAFDIYNDKVAGFAKFLKTNCCRFQEKDE<br>DDNLIDSYFVVKRHTFSNYQHEETIYNLLKDCPAVAKHDF<br>FKFRIDGDMVPHISRQRLTKYTMADLVYALRHFDEGNCDT<br>LKEILVTYNCCDDDYFNKKDWYDFVENPDILRVYANLGER<br>VRQALLKTVQFCDAMRNAGIVGVLTLDNQDLNGNWYDF<br>GDFIQTTPGSGVPVVDSYYSLLMPILTLTRALTAESHVDTD<br>LTKPYIKWDLLKYDFTEERLKLFDRYFKYWDQTYHPNCV<br>NCLDDRCILHCANFNVLFSTVFPPTSFGPLVRKIFVDGVPFV<br>VSTGYHFRELGVVHNQDVNLHSSRLSFKELLVYAADPAM<br>HAASGNLLLDKRTTCFSVAALTNNVAFQTVKPGNFNKDF<br>YDFAVSKGFFKEGSSVELKHFFFAQDGNAAISDYDYYRYN<br>LPTMCDIRQLLFVVEVVDKYFDCYDGGCINANQVIVNNLD<br>KSAGFPFNKWGKARLYYDSMSYEDQDALFAYTKRNVIPTI<br>TQMNLKYAISAKNRARTVAGVSICSTMTNRQFHQKLLKSI<br>AATRGATVVIGTSKFYGGWHNMLKTVYSDVENPHLMGW<br>DYPKCDRAMPNMLRIMASLVLARKHTTCCSLSHRFYRLA<br>NECAQVLSEMVMCGGSLYVKPGGTSSGDATTAYANSVFN<br>ICQAVTANVNALLSTDGNKIADKYVRNLQHRLYECLYRNR<br>DVDTDFVNEFYAYLRKHFSMMILSDDAVVCFNSTYASQG<br>LVASIKNFKSVLYYQNNVFMSEAKCWTETDLTKGPHEFCS<br>QHTMLVKQGDDYVYLPYPDPSRILGAGCFVDDIVKTDGTL<br>MIERFVSLAIDAYPLTKHPNQEYADVFHLYLQYIRKLHDEL<br>TGHMLDMYSVMLTNDNTSRYWEPEFYEAMYTPHTVLQ | 108602 |
| nsp12-<br>Twins-<br>Strep | MSADAQSFLNRVCGVSAARLTPCGTGTSTDVVYRAFDIYN<br>DKVAGFAKFLKTNCCRFQEKDEDDNLIDSYFVVKRHTFSN<br>YQHEETIYNLLKDCPAVAKHDFFKFRIDGDMVPHISRQRLT<br>KYTMADLVYALRHFDEGNCDTLKEILVTYNCCDDDYFNK<br>KDWYDFVENPDILRVYANLGERVRQALLKTVQFCDAMRN<br>AGIVGVLTLDNQDLNGNWYDFGDFIQTTPGSGVPVVDSYY<br>SLLMPILTLTRALTAESHVDTDLTKPYIKWDLLKYDFTEER | 110117 |

| | | |
|---|---|---|
| | LKLFDRYFKYWDQTYHPNCVNCLDDRCILHCANFNVLFST VFPPTSFGPLVRKIFVDGVPFVVSTGYHFRELGVVHNQDV NLHSSRLSFKELLVYAADPAMHAASGNLLLDKRTTCFSVA ALTNNVAFQTVKPGNFNKDFYDFAVSKGFFKEGSSVELKH FFFAQDGNAAISDYDYYRYNLPTMCDIRQLLFVVEVVDKY FDCYDGGCINANQVIVNNLDKSAGFPFNKWGKARLYYDS MSYEDQDALFAYTKRNVIPTITQMNLKYAISAKNRARTVA GVSICSTMTNRQFHQKLLKSIAATRGATVVIGTSKFYGGW HNMLKTVYSDVENPHLMGWDYPKCDRAMPNMLRIMASL VLARKHTTCCSLSHRFYRLANECAQVLSEMVMCGGSLYV KPGGTSSGDATTAYANSVFNICQAVTANVNALLSTDGNKI ADKYVRNLQHRLYECLYRNRDVDTDFVNEFYAYLRKHFS MMILSDDAVVCFNSTYASQGLVASIKNFKSVLYYQNNVF MSEAKCWTETDLTKGPHEFCSQHTMLVKQGDDYVYLPYP DPSRILGAGCFVDDIVKTDGTLMIERFVSLAIDAYPLTKHP NQEYADVFHLYLQYIRKLHDELTGHMLDMYSVMLTNDNT SRYWEPEFYEAMYTPHTVLQLEGGGGWSHPQFEKGGGSG GGSGGGSWSHPQFEK | |
| N-protein | MSDNGPQNQRNAPRITFGGPSDSTGSNQNGERSGARSKQR RPQGLPNNTASWFTALTQHGKEDLKFPRGQGVPINTNSSP DDQIGYYRRATRRIRGGDGKMKDLSPRWYFYYLGTGPEA GLPYGANKDGIIWVATEGALNTPKDHIGTRNPANNAAIVL QLPQGTTLPKGFYAEGSRGGSQASSRSSSRSRNSSRNSTPG SSRGTSPARMAGNGGDAALALLLLDRLNQLESKMSGKGQ QQQGQTVTKKSAAEASKKPRQKRTATKAYNVTQAFGRRG PEQTQGNFGDQELIRQGTDYKHWPQIAQFAPSASAFFGMS RIGMEVTPSGTWLTYTGAIKLDDKDPNFKDQVILLNKHIDA YKTFPPTEPKKDKKKKADETQALPQRQKKQQTVTLLPAAD LDDFSKQLQQSMSSADSTQALEGGGGWSHPQFEKGGGSG GGSGGGSWSHPQFEK | 48951 |
| pUL15 | MSYYHHHHHHHHDYDIPTTENLYFQGQQLASDVQQYLER LEKQRQLKVGADEASAGLTMGGDALRVPFLDFATATPKR HQTVVPGVGTLHDCCEHSPLFSAVARRLLFNSLVPAQLKG RDFGGDHTAKLEFLAPELVRAVARLRFKECAPADVVPQRN AYYSVLNTFQALHRSEAFRQLVHFVRDFAQLLKTSFRASS LTETTGPPKKRAKVDVATHGRTYGTLELFQKMILMHATYF LAAVLLGDHAEQVNTFLRLVFEIPLFSDAAVRHFRQRATV FLVPRRHGKTWFLVPLIALSLASFRGIKIGYTAHIRKATEPV FEEIDACLRGWFGSARVDHVKGETISFSFPDGSRSTIVFASS HNTNGIRGQDFNLLFVDEANFIRPDAVQTIMGFLNQANCKI IFVSSTNTGKASTSFLYNLRGAADELLNVVTYICDDHMPRV VTHTNATACSCYILNKPVFITMDGAVRRTADLFLADSFMQ EIIGGQARETGDDRPVLTKSAGERFLLYRPSTTTNSGLMAP DLYVYVDPAFTANTRASGTGVAVVGRYRDDYIIFALEHFF LRALTGSAPADIARCVVHSLTQVLALHPGAFRGVRVAVEG NSSQDSAVAIATHVHTEMHRLLASEGADAGSGPELLFYHC EPPGSAVLYPFFLLNKQKTPAFEHFIKKFNSGGVMASQEIV SATVRLQTDPVEYLLEQLNNLTETVSPNTDVRTYSGKRNG ASDDLMVAVIMAIYLAAQAGPPHTFAPITRVS | 83887 |
| pUL28 | MGAAPVSEPTVARQKLLALLGQVQTYVFQIELLRRCDPHI GRGKLPQLKLNALQVRALRRRLRPGLEAQAGAFLTPLSVT LELLLEYAWREGERLLGSLETFATAGDVAAFFTETMGLAR PCPYHQRVRLDTYGGTVHMELCFLHDVENFLKQLNYCHLI TPSRGATAALERVREFMVGAVGSGLIVPPELSDPSHPCAVC FEELCVTANQGATIARRLADRICNHVTQQAQVRLDANELR RYLPHAAGLSDADRARALSVLDHALARTAGGDGQPHPSP | 85635 |

| | | |
|---|---|---|
| | ENDSVRKEADALLEAHDVFQATTPGLYAISELRFWLASGD RAGQTTMDAFASNLTALARRELQQETAAVAVELALFGRR AEHFDRAFGSHLAALDMVDALIIGGQATSPDDQIEALIRAC YDHHLTTPLLRRLVSPEQCDEEALRRVLARMGAGGAADA PKGGAGPDDDGDRVAVEEGARGLGAPGGGGEDEDRRRGP GGQGPETWGDIATQAAADVRERRRLYADRLTKRSLASLG RCVREQRGELEKMLRVSVHGEVLPATFAAVANGFAARAR FCALTAGAGTVIDNRSAPGVFDAHRFMRASLLRHQVDPAL LPSITHRFFELVNGPLFDHSTHSFAQPPNTALYYSVENVGLL PHLKEELARFIMGAGGSGADWAVSEFQRFYCFDGISGITPT QRAAWRYIRELIIATTLFASVYRCGELELRRPDCSRPTSEGR YRYPPGVYLTYDSDCPLVAIVESAPDGCIGPRSVVVYDRD VFSILYSVLQHLAPRLPDGGHDGPP | |
| pUL33 | MGREGRTRQRTLRDTIPDCALRSQTLESLDARYVSRDGAH DAAVWFEDMTPAELEVVFPTTDAKLNYLSRTQRLASLLTY AGPIKAPDDAAAPQTPDTACVHGELLARKRERFAAVINRF LDLHQILRG | 14451 |
| pUL32 monomer | MSYYWSHPQFEKDYDIPTTENLYFQGATSPPGVLASVAVC EESPGSSWKAGAFERAYVAFDPSLLALNEALCAELLTASH VIGVPPVGTLDEDVAADVVTAPSRARGGAGDGGGSGGRG GPRNPPPDPCGEGLLDTGPFSAAAIDTFALDRPCLVCRTIEL YKQTYRLSPQWVADYAFLCAKCLGAPHCAASIFVAAFEFV YVMDRHFLRTKKATLVGSFARFALTINDIHRHFFLHCCFRT DGGVPGRHAQKQPKPSPSPGAAKVQYSNYSFLAQSATRAL IGTLASGGEEGAGSAAGSGTQPSLTTALMNWKDCARLLDC TEGRRGGGDSCCTRAAARNGEFETVAGDREPEESPDTWA YADLVLLLLAGTPAVWESGPQLRAAAEARRATVRQSWEA HRGARTRDVAPRFAQFTEPDAQPDLDLGPLMATVLKHGR GRGRTGGECLLCNLLLVRAYWLALRRLRASVVRYSENNT SLFDCIVPVVDQLEADPETQPGDGGRFVSLLRAAGPEAIFK HMFCDPMCAITEMEVDPWVLFGHPPATHPDELLLHKAKL ACGNEFEGRVCIALRALIYTFKTYQVFVPKPTALATFVREA GALLRRHSISLLSLEHTLCTYV* | 67061 |
| pORF68 | MWSHPQFEKGGGSGGGSGGGSWSHPQFEKGAAMFVPWQ LGTITRHRDELQKLLAASLLPEHPEESLGNPIMTQIHQSLQP SSPCRVCQLLFSLVRDSSTPMGFFEDYACLCFFCLYAPHCW TSTMAAAADLCEIMHLHFPEEEATYGLFGPGRLMGIDLQL HFFVQKCFKTTAAEKILGISNLQFLKSEFIRGMLTGTITCNF CFKTSWPRTDKEEATGPTPCCQITDTTTAPASGIPELARATF CGASRPTKPSLLPALIDIWSTSSELLDEPRPRLIASDMSELKS VVASHDPFFSPPLQADTSQGPCLMHPTLGLRYKNGTASVC LLCECLAAHPEAPKALQTLQCEVMGHIENNVKLVDRIAFV LDNPFAMPYVSDPLLRELIRGCTPQEIHKHLFCDPLCALNA KVVSEDVLFRLPREQEYKKLRASAAAGQLLDANTLFDCEV VQTLVFLFKGLQNARVGKTTSLDIIRELTAQLKRHRLDLAH PSQTSHLYA | 55097 |
